# A comparative transcriptomic analysis of mouse demyelination models and Multiple Sclerosis lesions

**DOI:** 10.1101/2025.03.24.645058

**Authors:** Erin L Aboelnour, Veronica R Vanoverbeke, Elizabeth A Maupin, Madelyn M Hatfield, Katrina L Adams

## Abstract

Demyelinating diseases are debilitating conditions characterized by loss of the myelin sheaths, ultimately leading to neurodegeneration. Toxicity models are among the most commonly used mouse models to induce demyelination; however, it remains unclear whether different demyelination models elicit distinct glial responses, and how comparable these changes are to human diseases like Multiple Sclerosis (MS). To address this gap, we integrated new and published single cell transcriptomic data of the subcortical white matter from lysophosphatidylcholine (LPC) and cuprizone (CPZ) toxicity models, and compared them to an existing human MS dataset. We find that CPZ demyelination induces a distinct oligodendrocyte (OL) state, enriched for cell stress pathways and C*dkn1a* and *Nupr1* expression, that resembles disease-associated OLs in MS lesions. However, both LPC and CPZ converge at a similar remyelination OL state that expresses immune-related genes, such as *Socs3* and *B2m*. Analysis of microglia reveals a largely conserved activation of microglia in both models, but LPC demyelination induces a stronger response, with recruitment of perivascular macrophages, that persists longer during remyelination. Interestingly, both mouse models do not recapitulate the heterogeneity of microglia across different MS lesion types. Overall, this comparative analysis uncovers specific gene expression differences between mouse demyelination models and human MS lesions, providing a foundation for using the animal models effectively to advance remyelination therapies.

## Introduction

Demyelination, the loss of myelin sheaths around neuronal axons, is a major contributor to neurodegenerative disease, such as Multiple Sclerosis (MS). Demyelination is spontaneously repaired in the young, healthy central nervous system (CNS), and is sufficient to provide neuroprotection and slow disease progression^1,2^. However, evidence of remyelination is highly variable across MS patients, and there are no approved treatments that promote remyelination. Therefore, there is an urgent need to understand the biology underlying remyelination. Remyelination is a complex, multicellular process that is most frequently studied using toxicity-induced demyelination in mice, due to the predictable timeline of damage and repair. The most widely used toxicity models to induce demyelination use lysolecithin (LPC) or cuprizone (CPZ)^3,4^. LPC is a detergent that creates a focal lesion that undergoes repair through reproducible phases of demyelination, increased reactive gliosis, and differentiation of oligodendrocyte precursor cells (OPCs) into oligodendrocytes (OLs) within approximately four weeks. CPZ is a copper chelator that, when fed to mice for four to six weeks, induces OL cell death throughout the grey and white matter (GM and WM, respectively) of the brain. After CPZ withdrawal, the brain undergoes spontaneous OPC differentiation and remyelination within two to three weeks^5,6^. These two toxicity models are often used interchangeably by researchers, with a particular method selected because of experimental design constraints rather than scientific rationale. However, it remains unclear whether LPC and CPZ induce similar molecular changes and cellular states.

The advancement of single cell (sc) and single nuclei (sn) RNA sequencing (sc/sn-RNA-seq) and supporting bioinformatic tools has enabled unbiased characterization of the molecular responses across neuronal and glial populations from a multitude of CNS pathologies ^7^. Previous studies have characterized the response to LPC and CPZ treatment using scRNA-seq ^8–10^. In particular, Pandey et al. described three distinct disease-associated oligodendrocyte (DAO) states that are similar in gene expression between mouse demyelination and Alzheimer’s disease models; however, they did not directly compare the effects of LPC and CPZ treatments ^9^. Likewise, snRNA-seq analysis of post-mortem human brain samples has provided valuable insight into the cellular and molecular pathology of MS ^11–14^. MS lesions, particularly the rims of chronic active lesions, contain stressed OLs, reactive astrocytes, and activated phagocytosing microglia/macrophages ^11,15,16^. While mouse models have been used to identify altered signaling pathways that appear conserved in MS lesions, a direct comparison of mouse toxicity-induced demyelination models to human MS lesion transcriptomics across multiple cell types has not been performed ^9,17^. A greater understanding of the similarities and differences between mouse models and MS lesions is critical for effectively using these models to identify and test new therapeutic targets.

In this study, we directly compare the effects of LPC- and CPZ-induced demyelination in subcortical WM. We integrate new and existing sc/snRNA-seq datasets of the demyelinated and remyelinated corpus callosum from LPC-injected and CPZ-fed mice ^8,9^. We also reanalyze a publicly available snRNA-seq dataset of the largest cohort of MS lesions to date. We first analyzed gene expression changes within the OL lineage and found that CPZ demyelination induces a strongly dysregulated disease-associated OL (DAO) state, marked by expression of *Cdkn1a* and *Nupr1*, that is not observed in LPC treatment. However, both LPC and CPZ converge on a similar remyelination DAO state characterized by upregulation of immune-response genes, such as *Socs3* and *B2m*. Importantly, we find that *CDKN1A* and *SOCS3* are also upregulated in human DAOs within MS lesions, and CPZ-demyelination better recapitulates the human DAO state, as evidenced by shared upregulated stress pathways and downregulation of myelin maintenance genes. We then analyzed gene expression changes within microglia and macrophages and found an opposite trend. LPC- and CPZ-induced demyelination elicit a conserved transcriptomic response in microglia, but LPC demyelination induces a stronger and more prolonged activation of disease-associated microglia/macrophages (DAMs). Interestingly, microglia in human MS lesions display more heterogeneity than the mouse models, with distinct DAM transcriptional states enriched across different MS lesion types. We identify a conserved group of DAM genes across species, but notably find that DAMs in remyelinated MS lesions enrich additional pathways similar to mouse microglia, including hypoxia and glycolysis.

## Results

### Compiling single cell/nuclei transcriptomic datasets of mouse demyelination models and MS patient samples

To compare the effects of LPC- and CPZ-induced demyelination on different WM cells, we performed snRNA-seq of the adult mouse corpus callosum, and integrated our data with two additional published scRNA-seq studies of the corpus callosum following LPC- or CPZ-induced demyelination ^8,9^. Together, these datasets cover a time course from baseline (homeostatic conditions) through demyelination and subsequent remyelination (Figure 1a, Supplementary Data 1). We obtained 112,017 high-quality filtered cells/nuclei, and the fully integrated UMAP displayed strong overlap between datasets and transcriptome types while maintaining clear cell-type separation (Figure 1b, Supplementary Figure 1a-c). Using expression of known marker genes, we annotated clusters corresponding to OL lineage cells, (OPCs, committed oligodendrocyte precursors (COPs), and mature OLs (MOLs)), microglia, perivascular macrophages (PVMs), astrocytes, neural progenitor cells (NPCs), ependymal cells, endothelial cells, vascular cells, pericytes, T cells, and three clusters undergoing cell division (OPCs, microglia, and NPCs) (Figure 1c, Supplementary Figure 1d-e). MOLs and microglia make up the majority of the data (Figure 1c).

**Figure 1:**
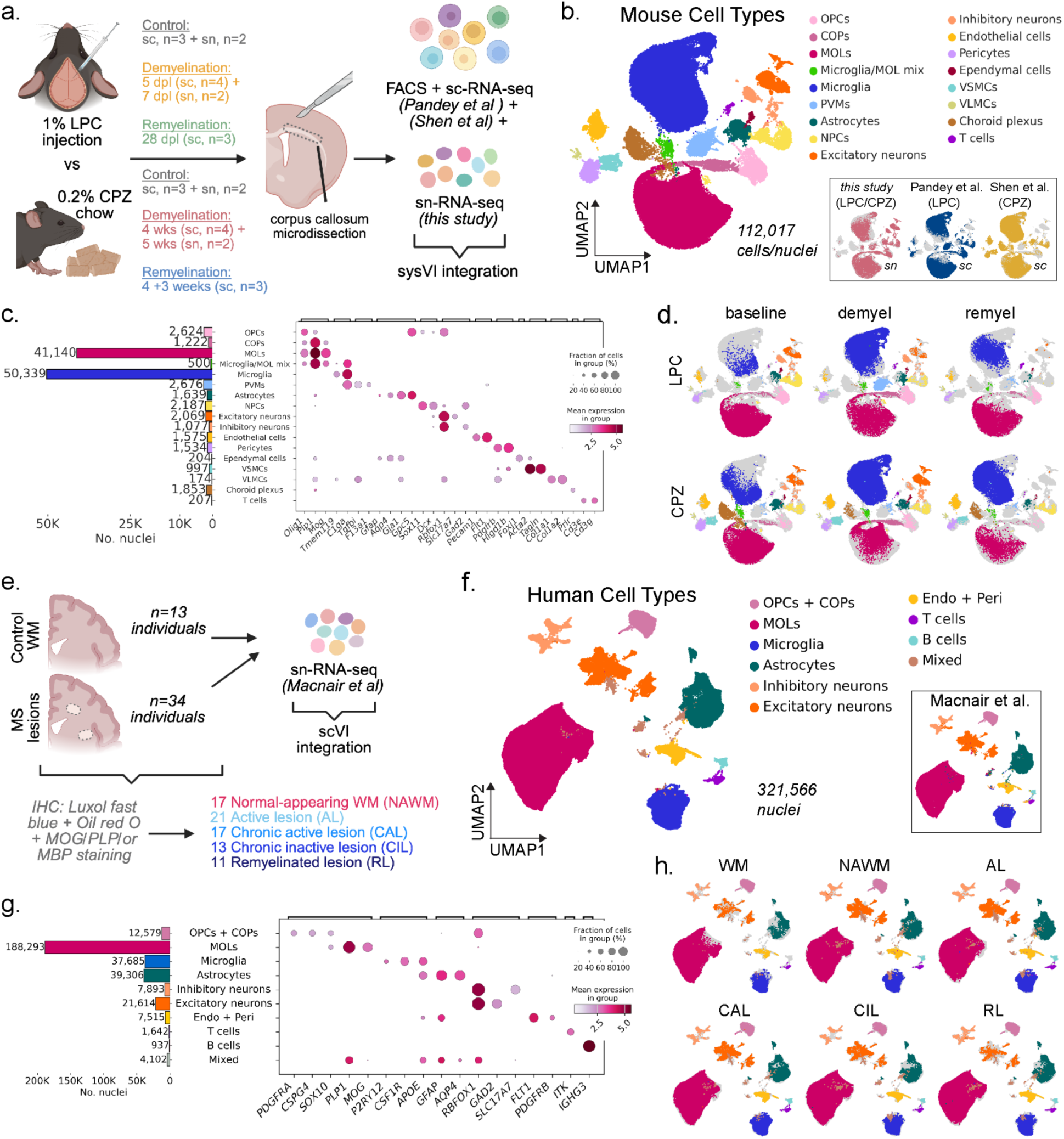
Single cell/nuclei RNA-seq datasets from mouse demyelination models and MS tissue. a. Schematic depicting mouse sc/snRNA-seq datasets included in this study. Adult corpus callosum tissue was microdissected from LPC- and cuprizone (CPZ)-induced demyelination/remyelination models, along with untreated controls. Single-cell suspensions were generated by enzymatic dissociation followed by FACS sorting. Replicate numbers for each condition and timepoint are indicated. Datasets were integrated using sysVI. Schematics created with BioRender. b. UMAP of the integrated mouse dataset annotated by cell types. Inset UMAPs colored by dataset of origin (publication source), labeled with demyelination treatment and library type. (OPCs, oligodendrocyte progenitor cells; COPs, committed oligodendrocyte progenitors; MOLs, mature oligodendrocytes; PVMs, perivascular macrophages; NPCs, neural precursor cells; VSMCs, vascular smooth muscle cells; VLMCs, vascular and leptomeningeal cells). c. Cell-type composition across the mouse dataset and marker validation. The bar chart shows total cell counts per annotated cell type. The corresponding dot plot demonstrates expression of canonical marker genes, where dot color represents mean expression and dot size indicates the proportion of cells expressing the gene. d. UMAPs showing the distribution of cell types across treatments (LPC or CPZ) and experimental stages (baseline, demyelination, remyelination). e. Overview of the human snRNA-seq dataset. White matter control samples (n = 13) and multiple sclerosis lesions (79 lesions from 34 patients) were integrated using scVI. Lesions were classified as WM, NAWM, active (AL), chronic active (CAL), chronic inactive (CIL), or remyelinated (RL) lesions based on histopathology. Schematics created with BioRender. f. UMAP of the integrated human dataset annotated by major cell types. Inset shows concordance with cell-type annotations from the original publication (Endo + Peri, endothelial cells and pericytes). g. Cell-type composition and marker validation for the human dataset, displayed as a bar chart and corresponding marker gene dot plot. h. UMAPs colored by cell-type annotation plotted for each lesion type.

Plotting the UMAPs separated by treatment and time point, we observed changes in cell type abundances and states in both LPC and CPZ, with primarily a reduction in MOLs and an increase in microglia at demyelination timepoints, as expected (Figure 1d, Supplementary Figure 1f-g). Although astrocytes are known to respond robustly in both demyelination models ^18,19^, we did not obtain sufficient numbers of astrocytes for robust downstream analysis. We also observed that PVMs were specifically enriched in the LPC model, showing distinctions at the cellular level between LPC- and CPZ-induced response. Using scCODA to statistically test these changes in cellular composition, we found PVMs and microglia increased in abundance, and MOLs decreased upon LPC demyelination (Supplementary Figure 1g). At the LPC remyelination timepoint, microglia remained elevated, with also increased T cells and MOLs, and modestly decreased Dcx^+^ NPCs (Supplementary Figure 1g) ^20^. Microglia also significantly increased in abundance in CPZ de- and remyelination, with only MOLs decreased in abundance at CPZ demyelination (Supplementary Figure 1g).

To generate a comparable human dataset to compare with the mouse demyelination models, we also reanalyzed a recently published snRNA-seq MS dataset, extracting only the WM samples from their atlas ^14^. In total, the human dataset includes 13 control WM samples from non-MS patients, 17 normal-appearing white matter (NAWM) samples, 21 active lesions (ALs), 17 chronic active lesions (CALs), 13 chronic inactive lesions (CILs), and 11 remyelinated lesions (RLs) from a total of 47 individuals (Figure 1e, Supplementary Data 2). We found that there was no statistically significant difference in the age, sex (25 female, 22 male), postmortem interval (PMI), or counts per nucleus between the control and MS individuals (Supplementary Figure 2a-c). Multiple lesions were collected from many individuals, totalling 92 samples, and most of the lesion samples are from patients diagnosed with more progressive forms of the disease – secondary progressive MS (SPMS) and primary progressive MS (PPMS) (Supplementary Figure 2d, and Supplementary Figure 2e-g for additional quality control).

After quality filtering, we integrated 321,565 nuclei and annotated the human cell types using canonical markers, which showed highly similar results to the previously published report (Figure 1f-g, Supplementary Figure 3a-c). Similar to the mouse dataset, this human dataset comprises mostly glial cells, with the majority coming from MOLs, as well as good representation of microglia and astrocytes. Plotting the UMAPs by lesion type revealed shifts in microglia, astrocyte, and MOL states across MS lesions and patient diagnoses (Figure 1h, Supplementary Figure 3d-e). Peripheral immune cells, including T and B cells, were predominantly detected in MS lesion samples, consistent with their known pathological roles in MS. scCODA analysis also showed a significant decrease in MOLs in ALs and CALs (Supplementary Figure 3d). Overall, these mouse demyelination and human MS datasets recapitulate expected cellular changes, allowing for detailed comparative transcriptomic analyses between mouse models and across species.

### Cuprizone demyelination induces a distinct DAO state not present in LPC demyelination

To understand how specific glial populations respond to LPC versus CPZ demyelination, we first focused on OL lineage cells. We subclustered and reannotated the OL lineage cells in our integrated mouse sn/scRNA-seq dataset (Figure 1b) using known markers for OPCs, COPs, newly formed OLs (NFOLs), myelin-forming OLs (MFOLs), and MOLs (Figure 2a,b, Supplementary Figure 4a-c). PAGA connectivity revealed a clear developmental progression of progenitors into mature states (Fig 2a, inset). We defined four homeostatic mouse MOL states (MOL_A-D), which all expressed mature markers (*Plp1*, *Mog*, *Mag, Opalin*), along with cluster-specific markers (MOL_A: *Il33^low^*; MOL_B: *Il33*^high^ and *Ptgds*^high^; MOL_C: increased expression of *Selenop*, *Spock3*, *Anln;* and MOL_D: *Klk6*, *S100b*, *Anxa5),* which have been defined in previous studies (Figure 2b) ^21,22^. MOL_B and MOL_C constituted the majority of adult mouse corpus callosum OLs, accounting for 75-80% of MOLs at baseline, consistent with a previous report of low *Klk6* expression in the adult mouse corpus callosum ^23^. Additionally, we identified three MOL subclusters that express genes associated with previously described DAO states, including *Serpina3n*, *C4b* and *H2-D1* (MOL_E and F) and *Cdkn1a*, *Ddit3* and *Nupr1* (MOL_G) (Figure 2b, Supplementary Figure 4c) ^9,10,17,22^.

**Figure 2:**
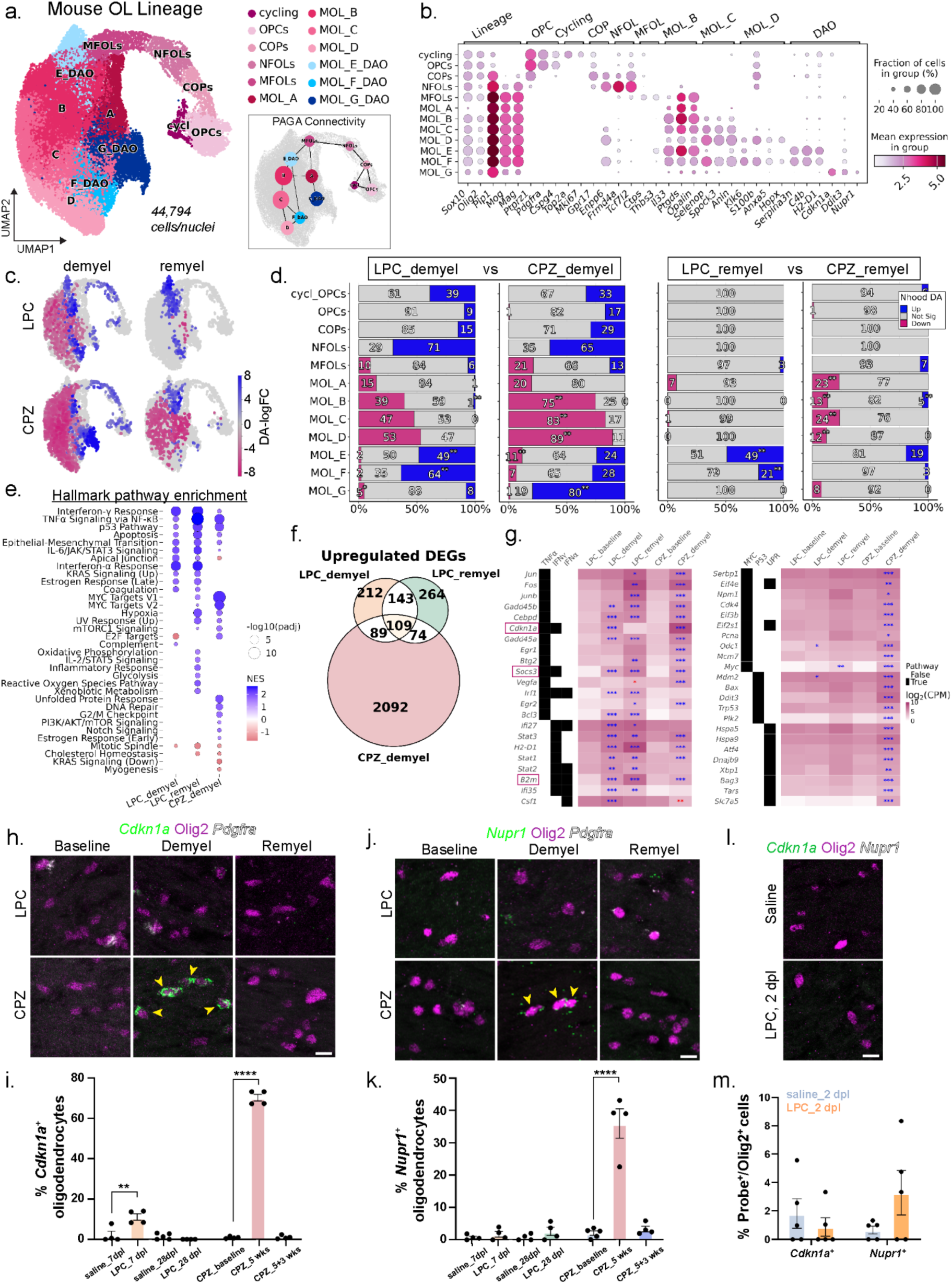
CPZ induces a distinct transcriptional state in oligodendrocytes during demyelination. a. UMAP embedding of the subset and reclustered mouse oligodendrocyte lineage cells. The inset UMAP shows PAGA connectivity with edge weights > 0.4 between annotated cell types. (OPCs, oligodendrocyte precursor cells; COPs, committed oligodendrocyte precursors; NFOLs, newly formed oligodendrocytes; MFOLs, myelin-forming oligodendrocytes; DAOs, disease-associated oligodendrocytes). b. Dot plot of lineage- and cluster-specific marker genes across oligodendrocyte lineage cells. c. Differential abundance (DA) of neighborhoods (nhoods) superimposed on the UMAP embedding. Each node represents a nhood and is colored by the log fold-change (logFC) relative to baseline; non-significant nhoods are shown in grey (spatial FDR < 0.1). d. Bar plot showing the proportion of non-significant (grey), increased (blue), and decreased (magenta) nhoods, as defined in panel c, for each cell type annotated in panel a. Significance annotations (* p < 0.05, ** p < 0.01) indicate results from cell-type-specific enrichment tests comparing LPC vs. CPZ at demyelination (left) and remyelination (right) using Fisher’s exact or chi-squared tests. Exact p-values are provided in Supplementary Table 3. e. Dot plot showing Hallmark pathway enrichment in the indicated condition. Dot color reflects the normalized enrichment score (NES) and dot size indicates –log10(adjusted p-value). f. Venn diagram illustrating shared and condition-specific significantly upregulated genes (logFC ≥ 1.5, adjusted p < 0.05) derived from pseudobulk MOLs in LPC or CPZ during demyelination and LPC during remyelination. g. Heatmap showing cytokine-associated gene expression in LPC and CPZ (left) and CPZ-specific enrichment of Myc, p53, and UPR-associated genes (right). Heatmaps display average log₂-transformed CPM expression. Gene–pathway assignments are derived from Hallmark gene sets, shown in the annotation panel (black tiles indicate membership). Genes are hierarchically ordered first by pathway and then by mean expression. Blue asterisks indicate significantly upregulated genes and red asterisks indicate significantly downregulated genes (* p < 0.05, ** p < 0.01, *** p < 0.001). Full results are provided in Supplementary Table 4. (TNFα, tumor necrosis factor alpha; IFNγ, interferon gamma; IFNα, interferon alpha). h. Representative RNAscope images of mouse corpus callosum showing *Cdkn1a* expression. Yellow arrows show *Cdkn1a^+^* oligodendrocytes. LPC timepoints: baseline = saline at 7 dpl, demyelination = LPC at 7 dpl, remyelination = LPC at 28 dpl. CPZ timepoints: baseline = normal diet, demyelination = 5 weeks CPZ, remyelination = 5 weeks CPZ + 3 weeks recovery. (Scale bar = 10 μm). i. Bar chart showing the percentage of *Cdkn1a*^+^ oligodendrocytes (*Cdkn1a*^+^Olig2^+^*Pdgfra*^-^ / Olig2^+^*Pdgfra*^-^). Bars show group means, dots show individual animals, and error bars represent the standard error of the mean (SEM). Statistical significance was determined by one-way ANOVA with Tukey’s test (\*\**p* = 0.0011, \*\*\*\**p* < 0.0001). j. Representative RNAscope images from the mouse corpus callosum showing *Nupr1* expression. Yellow arrows show *Nupr1^+^* oligodendrocytes. Timepoints as in panel h. (Scale bar = 10 μm). k. Bar chart showing the percentage of *Nupr1*^+^ oligodendrocytes (*Nupr1*^+^Olig2^+^*Pdgfra*^-^ / Olig2^+^*Pdgfra*^-^), as presented in panel i (\*\*\*\**p* < 0.0001). l. Representative RNAscope images showing a lack of *Cdkn1a* and *Nupr1* expression in Olig2^+^ cells at 2 dpl after saline (top) or LPC injection (bottom). (Scale bar = 10 μm). m. Bar chart showing the percent of *Cdkn1a*^+^ (left) and *Nupr1*^+^ (right) oligodendrocyte lineage cells (*Probe^+^*Olig2^+^/Olig2^+^) at 2 dpl in saline or LPC injected tissue. Bars show group means, dots show individual animals, and error bars indicate SEM.

We hypothesized that the DAO states would be enriched in the demyelination timepoints. To test this, we quantified the differential abundance (DA) across overlapping cell neighborhoods (nhoods) using Milo ^24^. As expected, both LPC and CPZ resulted in widespread MOL loss and moderate gains in progenitor cells at demyelination (Figure 2c, Supplementary Data 3). To determine if there were differences in composition between CPZ and LPC, we performed a separate statistical test comparing the distribution of nhoods by aggregating the counts of upregulated, downregulated, and non-significant nhoods by cell type and condition. We found that LPC-induced demyelination preferentially enriched DAO states MOL_E and F, whereas CPZ-induced demyelination resulted in an accumulation of the DAO MOL_G state (Figure 2d). Interestingly, LPC samples at remyelination showed near-complete recovery of homeostatic MOL populations, whereas CPZ samples exhibited persistent MOL depletion (Figure 2d).

To determine how LPC- and CPZ-induced demyelination alters MOL gene expression, we next tested for differentially expressed genes (DEGs) using a pseudobulk approach, comparing LPC and CPZ de- and remyelination MOLs to their respective baseline controls ((Supplementary Data 4). CPZ remyelination samples exhibited high heterogeneity, resulting in high p-values and low confidence in DEG calculations, so it was omitted from this analysis. Overall, we found similar numbers of DEGs at the different LPC timepoints (ex: 902 and 1,044 DEGs at LPC de- and remyelination, respectively), but a much higher number of DEGs at CPZ demyelination (4,230 genes), indicating that CPZ induces a very dramatic shift in MOL gene expression (Supplementary Figure 4d). We then performed gene set enrichment analysis (GSEA) to determine what signaling pathways are enriched in MOLs across demyelination (Figure 2e). TNF⍺ signaling via NFκB and interferon (IFN) signaling were the primary pathways enriched across all conditions, supported by shared upregulation of genes *Gadd45a/b*, *Stat1/3*, *B2m*, *H2-D1*, and *Socs3* (Figure 2e-g, Supplementary Figure 4e). LPC de- and re-myelination MOL DEGs were especially enriched for IFNγ and IFN⍺ signaling, marked by upregulation of *Irf1*, *Ifi27*, *Ifi35*, and *Stat2.* In contrast, CPZ demyelination MOL DEGs were strongly enriched for Myc targets (*Eif4e*, *Eif4a1*, *Odc1*), DNA repair (*Tp53*, *Ddb1, Pcna*), and unfolded protein response (UPR) (*Atf4*, *Hspa5*), indicating increased protein synthesis and dysregulated stress responses (Figure 2e-g). Consistent with our findings, CPZ treatment has been shown to increase oxidative stress, elevate production of reactive oxygen species (ROS), and induce endoplasmic reticulum stress, triggering UPR and OL cell death ^25,26^.

To confirm that the CPZ-induced DAO signature is not observed with the LPC model, we performed RNAscope for two highly upregulated genes in the mouse MOL_G DAO cluster: *Cdkn1a* and *Nupr1*. *Cdkn1a* encodes for p21, a p53-dependent kinase that is induced in response to DNA damage. *Nupr1* encodes a transcriptional regulator involved in the oxidative stress response, DNA damage, and ferroptosis. As expected, we found significant upregulation of both genes in OLs upon CPZ demyelination, which was gone by remyelination (3 weeks following removal of CPZ from diet) (Figure 2h-k). Conversely, we did not observe a significant change in either *Cdkn1a* or *Nupr1* expression in OLs at LPC de- and remyelination (Figure 2h-k). Given that CPZ induces a more continuous and protracted OL loss, we wondered if a similar stress state is present at earlier stages of LPC-induced demyelination. However, analysis of WM OL lineage cells in LPC-injected brains at 2 days post lesion (dpl) revealed no detected expression of *Cdkn1a* or *Nupr1*, confirming that CPZ-induced stress mechanisms are distinct from those induced by LPC (Figure 2l,m).

### At remyelination stages, LPC and cuprizone converge onto a shared DAO state that upregulates immune-related genes

Our analysis of different MOL states across the timeline of demyelination and repair showed that the mouse MOL_E DAO cluster remains elevated in both LPC and CPZ models at remyelination stages. To verify the presence of this population in both models, we performed RNAscope for two genes highly expressed in the MOL_E DAO cluster, *Socs3* and *B2m.* Socs3 is a member of the suppressor of cytokine signaling (SOCS) family of proteins and negatively regulates immune responses and inflammation. *B2m* encodes the beta-2 microglobulin protein, which is the light chain subunit of the Class I Major Histocompatibility Complex (MHC-I) molecule. We observed a significant increase in the expression of both genes in WM OLs of LPC-injected and CPZ-fed mice, compared to their respective controls (Figure 3a-d). LPC maintained similar percentages of *Socs3*+ and *B2m*+ OLs from demyelination (7dpl) to remyelination (28dpl) timepoints, whereas the percentages gradually increased from CPZ demyelination (5 wks) to remyelination (5+3 wks) timepoints (Figure 3b,d). Notably, MFOLs showed a direct connection to MOL_E in our PAGA connectivity analysis (Figure 2a), altogether suggesting that OLs may undergo an altered maturation state following demyelination.

**Figure 3:**
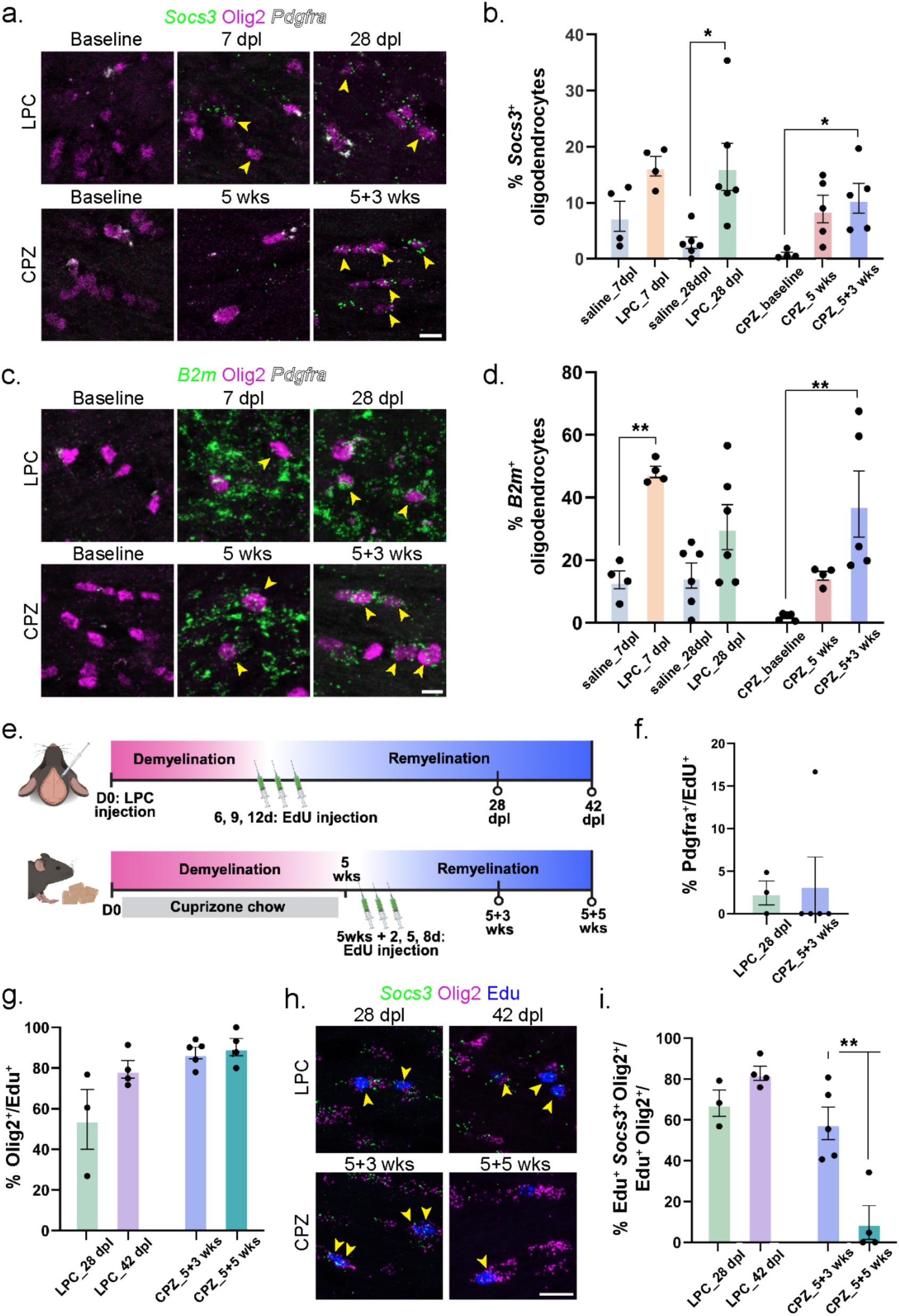
A convergent oligodendrocyte state expressing immune-related genes emerges during remyelination in both CPZ and LPC models. a. Representative RNAscope images of mouse corpus callosum showing *Socs3* expression. Yellow arrows indicate *Socs3*^+^ oligodendrocytes. LPC baseline = saline at 7 dpl, and CPZ baseline = normal diet. (Scale bar = 10 μm). b. Bar chart showing the percentage of *Socs3*^+^ oligodendrocytes (*Socs3*^+^Olig2^+^*Pdgfra*^-^ / Olig2^+^*Pdgfra*^-^). Bars show group means, dots show individual animals, and error bars represent the SEM. Statistical significance was assessed by one-way ANOVA with Tukey’s test (LPC: \**p* = 0.0125, CPZ: \**p* = 0.0273). c. Representative RNAscope images from the mouse corpus callosum showing *B2m* expression. Yellow arrows show *B2m^+^* oligodendrocytes. Timepoints are the same as panel a. (Scale bar = 10 μm). d. Bar chart showing the percentage of *B2m*^+^ oligodendrocytes (*B2m*^+^Olig2^+^*Pdgfra*^-^ / Olig2^+^*Pdgfra*^-^), as detailed in panel b. (LPC: \*\**p* = 0.0037, CPZ: \*\**p* = 0.0057). e. Schematic of EdU labeling experiment for LPC (top) and CPZ (bottom). For LPC, mice received saline or LPC injection at D0, followed by IP injection of EdU at 6, 9, and 12 dpl, and tissue was collected at 28 dpl (n = 3) or 42 dpl (n = 4). For CPZ, mice were fed CPZ chow for 5 weeks (wks), then returned to normal diet and injected with EdU at 2, 5, and 8 days post-switch, with tissue collected at 5 + 3 wks (n = 5) and 5 + 5 wks (n = 4). f. Bar chart showing the percentage of EdU^+^ OPCs (EdU^+^ Pdgfra^+^/EdU^+^) in LPC at 28 dpl and CPZ at 5+3 wks. Bars show group means, dots show individual animals, and error bars represent SEM. g. Representative RNAscope images from the mouse corpus callosum showing *Socs3* expression in LPC and CPZ treated mice. Yellow arrows indicate *Socs3^+^* EdU^+^ Olig2^+^ triple positive cells. (Scale bar = 10 μm). h. Bar chart showing the percentage of EdU^+^ cells that are oligodendrocyte lineage cells (EdU^+^ *Socs3*^+^ Olig2^+^/EdU^+^ Olig2^+^) in LPC- and CPZ-treated groups at the indicated timepoints. Bars represent group means, dots show individual animals, and error bars indicate SEM. i. Bar chart showing the percentage of newly-generated oligodendrocyte lineage cells that express *Socs3* (EdU^+^ *Socs3*^+^ Olig2^+^/EdU^+^ Olig2^+^). Bars show group means, dots show individual animals, and error bars indicate SEM. Statistical significance was assessed by an unpaired t-test with Welch’s correction (\**p* = 0.004121).

To determine if *Socs3* is expressed by newly generated MOLs during remyelination, we performed EdU labeling of proliferating progenitors at demyelination timepoints and collected the brains for analysis at two remyelination timepoints per model (Figure 3e). We confirmed that the majority of EdU-labeled cells express OL lineage marker Olig2, with a very small percentage expressing the OPC marker Pdgfra, indicating that the EdU protocol was successful in labeling newly-generated OLs (Figure 3f-h). We found that the majority of Olig2+EdU+ cells expressed *Socs3* in both LPC and CPZ demyelination (Figure 3i). Interestingly, *Socs3* expression remained high in these newly-generated OLs even at 42 days post LPC injection (5 weeks after our demyelination timepoint), but dropped significantly at 5 weeks removal from CPZ diet (Figure 3i), again suggesting slight differences in recovery timelines between the two models.

Overall, across LPC and CPZ demyelination, MOLs upregulate a shared IFN- and stress-associated gene program, including MHC-I antigen presentation genes (*H2-D1, H2-K1, B2m, Tap2, Nlrc5*), transcriptional mediators of cytokine signaling (*Stat1/2/3, Irf1/3/9*), and interferon-stimulated genes (*Ifi27, Rtp4, Zc3hav1*). This state is accompanied by upregulation of complement and glial stress markers (*Serpina3n, C4b, Gadd45a/b, Socs3*), and cytoskeletal remodeling factors (*Vim, Anxa2, Cd63*), suggesting that MOLs actively respond to inflammatory cues in a manner similar to that observed in experimental autoimmune encephalomyelitis (EAE). Previous studies have also found upregulation of immune-related genes within OL lineage cells in the EAE mouse model and post-mortem MS brain tissue ^11,17,27^. To see how gene expression changes in the LPC and CPZ toxicity models compare with those in the EAE model, we used a previously published DEG list generated from scRNA-seq of OL lineage cells isolated from the spinal cords of EAE mice (Supplementary Figure 5a,b, Supplementary Data 4)^27^. We examined the DEG overlap across the three models and found 27% and 22% of upregulated EAE DEGs were also upregulated in LPC and CPZ MOLs, respectively (Supplementary Figure 5c-e). Interestingly, CPZ and EAE shared a higher number of downregulated DEGs, compared to LPC and EAE (23% of EAE downregulated genes are shared with CPZ, only 7.5% are shared with LPC). Both EAE and CPZ MOLs downregulated metabolic and membrane maintenance genes (*Slc1a3, Abhd12, Gprc5b, Stard5*) and structural/adhesion factors (*Ncam1, Dclk1, Tspan15*), suggesting destabilization of the mature MOL phenotype and reduced metabolic resilience in both models. Interestingly, both EAE and CPZ MOLs also downregulated expression of epigenetic modifiers (*Dnmt3a*, *Hdac11*, *Tet1*), which may partially explain the larger changes in MOL gene expression in these models, compared to LPC.

Overall, the activation of TNF⍺, IFN, Myc, and UPR signaling pathways suggests that both LPC and CPZ induce OL stress, albeit through distinct mechanisms. Interestingly, CPZ induces a distinct demyelination-associated MOL state that converges on a similar remyelination fate as LPC, with elevated expression of immune-related genes. This highlights that immune OLs are not unique to the EAE model, thereby providing opportunities to explore how demyelination induces immune responses in OLs independent of infection or T cell activation.

### Cuprizone demyelination recapitulates the DAO state in MS lesions

The observation of distinct DAO states in the mouse models prompted us to next investigate whether these states resemble those found in MS lesions. To address this, we first subclustered and reannotated the human OL lineage cells from the snRNA-seq MS WM dataset in Figure 1 (Figure 4a, Supplementary Figure 6a). Using known marker gene expression, we identified human OPCs, COPs, and 4 MOL populations (MOL_A-D) (see Methods, Figure 4b, Supplementary Figure 6b). Human MOL_A is characterized by high *RBFOX1* and *SLC5A11* expression, while human MOL_B additionally expresses low levels of *CTNND2* and *PLXDC2*. Human MOL_C loses expression of MOL_A markers, further upregulates MOL_B markers, and expresses *OPALIN* and *LAMA2*. Human MOL_D is distinguished from MOL_C by increased expression of *FTL* and *CSPRP1*. Additionally, we identified clusters that express markers for DAOs (human MOL_E-G). Specifically, human MOL_E expresses high levels of *HSPH1*, *FOS*, and *JUN*, while MOL_F and MOL_G are marked by *CDKN1A*, *NEDD4L*, and *GADD45B* expression. We confirmed our DAO annotation using a DAO gene signature from a previous MS study, again showing that known DAO genes are enriched in MOL_E-F and OPC clusters (Supplementary Figure 6c) ^9^. PAGA connectivity analysis revealed a primary differentiation trajectory from OPCs to COPs, followed by MOL_A through B, C, and D, with additional connectivity between the DAO states (Supplementary Figure 6d).

**Figure 4:**
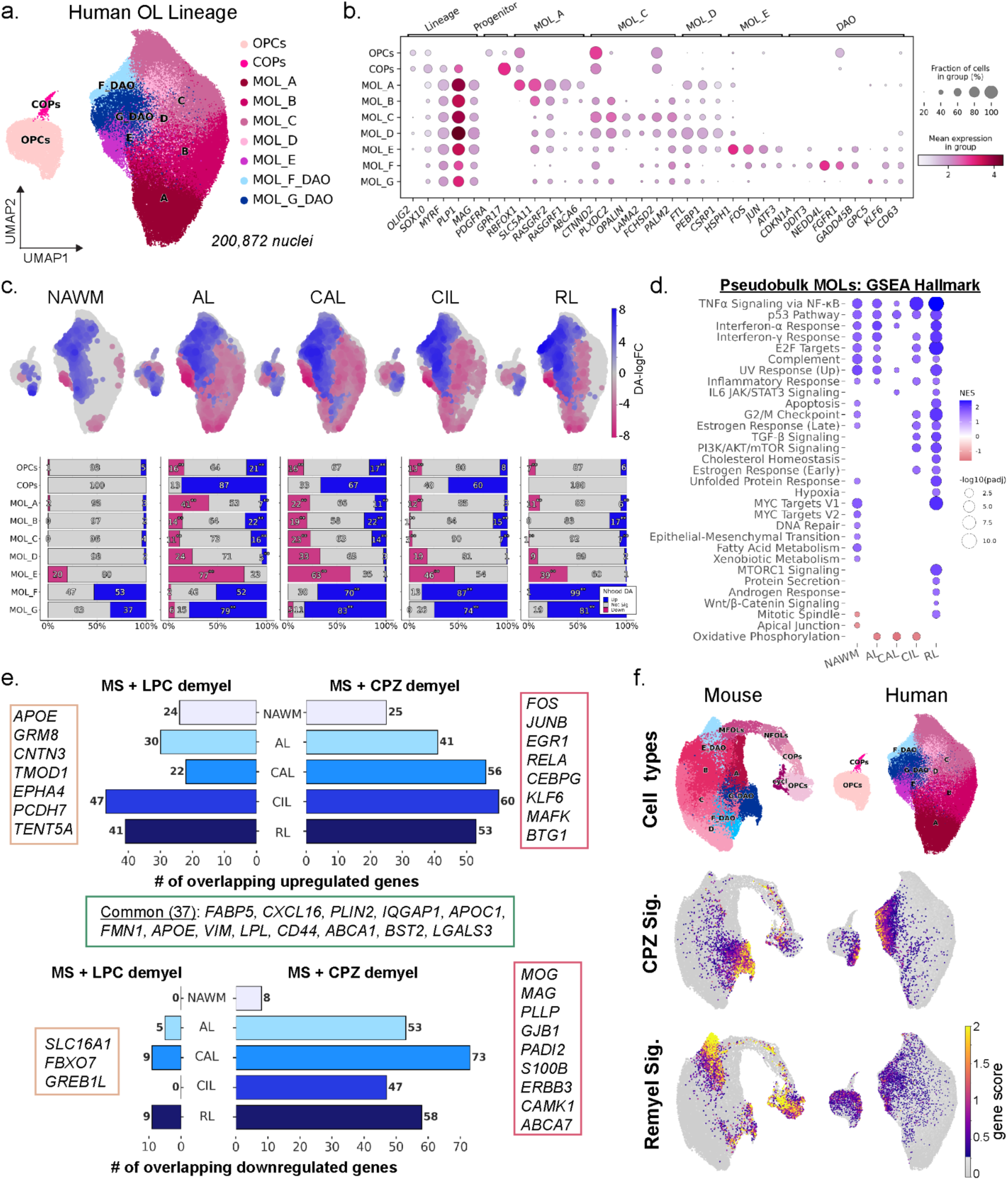
Mouse demyelination models share transcriptional phenotypes with disease-associated oligodendrocytes in human MS Lesions. a. UMAP embedding of the subset and reclustered human oligodendrocyte lineage cells. b. Dot plot of lineage- and cluster-specific marker genes across cell types. c. UMAP plots showing differential abundance (DA) neighborhoods (nhoods) assigned to annotated cell types for each lesion type compared with WM, computed using Milo. Blue indicates significantly increased nhoods, magenta indicates significantly decreased, and grey indicates non-significant (spatial FDR < 0.1). Bar plots (bottom) showing the proportion of non-significant (grey), increased (blue), and decreased (magenta) nhoods for each cell type. Significance annotations (* for *p* < 0.05 and ** for *p* < 0.01) indicate results from a separate statistical test assessing cell-type-specific enrichment using pairwise Fisher’s exact or chi-squared tests comparing the lesions to NAWM. Exact p-values are provided in Supplementary Table 5. d. Dot plot of Hallmark pathway enrichment across per lesion type. Dot color indicates normalized enrichment score (NES), and size reflects –log10(adjusted *p*). e. Bar charts showing overlaps of upregulated (top) and downregulated (bottom) MOL DEGs in LPC (left) and CPZ (right) at demyelination compared with MS lesion types. Genes shown in side panels occur in ≥ 1 shared comparison. The green box marks genes upregulated across both LPC and CPZ models and at least one MS lesion type. f. UMAP plots of mouse (left) and human (right) OL lineage cells and gene scores derived from CPZ-associated and remyelination-associated mouse DEGs.

To explore the distribution of these OL lineage states across the different MS lesions, we performed differential abundance testing using Milo (Supplementary Data 5). We found minimal changes in homeostatic MOL_A-D in NAWM compared to control WM, but variable alterations in homeostatic MOL_A-D in MS lesions, with the fewest changes observed in RLs (Figure 4c). All MS lesion types were strongly enriched for human DAO states MOL_F and G (Figure 4c). Interestingly, MOL_E, a comparatively small cluster, was highly enriched in two control WM samples (44% of cells), both from 85-year-old individuals (Supplementary Figure 6e). Human MOL_E shares similar gene expression changes to the DAO, suggesting that normal brain aging may induce a stressed state in MOLs.

We then performed pseudobulked differential gene expression testing for MOLs across the different MS lesion types, comparing each lesion to control WM (Supplementary Data 6). CAL MOLs had the largest number of DEGs (1,629 genes) and NAWM MOLs had the least (328 genes), with many DEGs shared across lesion types (Supplementary Figure 7a,b). Pathway enrichment analysis revealed upregulation of TNF⍺ signaling (*NFKB1A*, *BIRC3, CEBPD*), p53 pathway (*IER5, CDKN2A, CDKN2B*) and IFN⍺ response (*IRF7*) across multiple MS lesion types (Figure 4d), similar to what we observed with the mouse models. MOLs in CIL and RL lesions shared additional pathways, including TGFβ and PI3K/AKT/mTOR signaling (Figure 4d).

To compare the human and mouse datasets, we first used MetaNeighbor analysis to assess how well cell types correspond across the two species (Supplementary Figure 7c). Cell types from both species grouped into two main clusters, representing progenitors and MOLs. Within the MOL branch, we identified three subclusters, one containing DAOs and two containing homeostatic clusters, indicating shared cell identities between human and mouse. We then examined how closely the mouse toxicity models recapitulate the human DAO states by calculating the number of shared DEGs between LPC or CPZ and the different MS lesion types (Figure 4e, Supplementary Figure 7d). LPC demyelination MOLs shared 22-47 upregulated DEGs with MS MOLs, whereas CPZ demyelination MOLs shared 25-60 (Figure 4e). In particular, multiple mouse DAO genes were upregulated in CIL and/or RL lesions, including *SOCS3, CDKN1A, FOS, JUNB,* and *GADD45A/B (*Supplementary Figure 7f). *NUPR1* was also upregulated in CALs, but didn’t reach our significance threshold (log2FC = 1.48, FDR = 0.0703). Interestingly, *B2M* was not significantly upregulated by MOLs in any MS lesion type, however it has been shown to be upregulated in other MS datasets ^11–13^. When we examined the number of overlapping downregulated genes, we found that MOLs in CPZ demyelination shared more with MS lesions than MOLs in LPC de- and remyelination (Figure 4e). Many of these downregulated genes are involved in OL homeostasis and function, including myelin structural component genes *MOG*, *MAG*, and *PLLP*, myelin stability factor *ERBB3,* and lipid metabolic genes such as *ABCA7*. Downregulation of these genes suggests that DAOs in the CPZ model and in MS patients have a compromised ability to produce stable myelin.

To quantify transcriptional similarity, we calculated the overlap coefficient and Spearman correlation in a pairwise manner across mouse timepoints and human lesion conditions for upregulated and downregulated DEGs (Supplementary Fig. 7e, Supplementary Data 7). As expected, we observed strong similarity among the MS lesion states, reflecting a coherent DAO trajectory, with high Spearman correlations showing the logFC direction and magnitude is similar. CPZ showed a modest degree of overlap with MS lesions, particularly CIL (overlap coefficient = 0.41 for upregulated and 0.42 for downregulated genes), whereas LPC conditions exhibited much lower similarity, especially for downregulated genes (highest overlap coefficient = 0.06 between LPC demyelination and RL). The Spearman correlations were low in both LPC and CPZ comparisons, indicating that even where gene sets overlap, the direction and magnitude of fold-changes are not well aligned.

Finally, to examine if the different mouse DAO states resemble human DAOs in MS lesions, we generated gene signatures representing the CPZ-specific DAO state (mouse cluster MOL_G) and the remyelination DAO state (mouse cluster MOL_E). Interestingly, we see strong enrichment of the CPZ MOL_G gene signature within the human DAO clusters, indicating that CPZ demyelination models the human DAO state in MS lesions (Figure 4f). Genes in this state include stress response factors (*ATF4, DDIT3, NUPR1*), genes involved in injury response and hypoxia (*VEGFA, SERPINE1*), and pro-inflammatory genes (*CD44, LGALS3*). We also see enrichment of the remyelination MOL_E gene signature within the human DAO clusters, albeit to a lesser degree (Figure 4f). Genes in this state include negative regulators of immune signaling (*SOCS3, BCL3, TNFRSF1A, GADD45B/G*) and several components of ECM/lamina and membrane remodelers (*CD63, ANXA2, SNX10*). Altogether, these findings indicate that both mouse demyelination models recapitulate some key aspects of DAOs in MS lesions, such as increased TNF⍺ and IFN signaling. However, CPZ demyelination appears to more faithfully recapitulate the human DAO state in MS tissue. Finally, these results highlight *SOCS3* and *CDKN1A* as conserved DAO genes between mouse and human demyelination, warranting future studies of their functional roles in OL pathology.

### Mouse OPCs express DAO genes after demyelination and resemble OPCs in chronic inactive and remyelinated MS lesions

In addition to MOLs, we also examined OPC responses, given their critical role in remyelination. In addition to differentiating into new OLs, OPCs induce immune activation and participate in debris phagocytosis ^27–29^. To characterize their response, we analyzed gene expression changes in mouse OPCs at the LPC-induced demyelination timepoint (Supplementary Data 4). Unfortunately, few CPZ demyelination samples met our threshold for pseudobulking (at least 30 cells), so we could not robustly calculate OPC gene expression changes in the CPZ model. At LPC demyelination, we found 553 upregulated genes and 1,012 downregulated genes in mouse OPCs, with many genes and enriched pathways shared with DAO states in both LPC and CPZ demyelination. (Supplementary Figure 8a,b). Immune-related and IFN response genes were upregulated, including *Ifit1*, *Stat2*, *Socs3*, and *C4b*. Additionally, genes associated with cell cycle arrest and DNA damage response, such as *Gadd45b* and *Gadd45g*, were also upregulated. Interestingly, we also observe downregulation of genes such as *Mertk*, which could indicate that OPCs are suppressing their phagocytic function in favor of differentiation. Since our DAO gene signatures show enrichment within the OPC clusters (Figure 4f), we further examined the expression of DAO-associated genes in OPCs. Using RNAscope, we quantified the percentage of *Pdgfra*^+^Olig2^+^ OPCs that express *Cdkn1a*, *Nupr1*, *Socs3*, or *B2m* following LPC and CPZ demyelination. We found that *Cdkn1a* was significantly upregulated in OPCs at demyelination timepoints in both models, but was absent upon remyelination (Supplementary Figure 8c,d). Conversely, *Nupr1* expression did not significantly change in OPCs, although it did slightly increase at CPZ demyelination (Supplementary figure 8e). Both *Socs3* and *B2m* expression increased in OPCs across the two mouse models we found, remaining elevated at remyelination timepoints (Supplementary Figure 8f-h). These results indicate OPCs share a similar state across mouse models, with elevation of stress and immune-related genes that may impact OPC differentiation or survival.

We then similarly examined differential gene expression in human OPCs across the different MS lesion types (Supplementary Data 8). OPCs in CALs have the greatest number of DEGs (150 up and 231 downregulated), whereas OPCs in NAWM and ALs have the fewest changes in gene expression (Supplementary Figure 9a). Interestingly, there is a wide variety of GSEA pathways in OPCs across the different MS lesions, including downregulation of many pathways in CALs (cholesterol homeostasis, IFN signaling, fatty acid metabolism) and upregulation of pathways in RLs (hypoxia, glycolysis, and TNF⍺ signaling) (Supplementary Figure 9b,c). This suggests that OPCs exist in very different metabolic and transcriptomic states between different lesion types, which was confirmed by examining the number of shared DEGs across lesion types. Only 8 upregulated genes (*CASZ1, ARHGAP24, TRPM8, HELB, SULF1, NEURL1B, DPYD, TLE2*) and 10 downregulated genes (*COL9A3, CCND1, SERINC1, SNX22, PEX5L, CD82, FRMD4B, FKRP, AC113346.2, CIRBP*) were shared between OPCs in CALs and RLs (Supplementary Figure 9d). Overall, the highest number of shared DEGs was between CIL and RLs (38 upregulated, 16 downregulated), suggesting human OPCs are more similar in these lesions. Interestingly, human OPCs in CIL and/or RLs also upregulated several genes shared with mouse OPCs, including *SOCS3*, *TGFB2*, *GADD45B*, and *SNX10* (Supplementary Figure 9e). This is consistent with similar pathway enrichment in mouse OPCs and human OPCs in RLs (Myc targets, IL6/JAK/STAT, TNF⍺ signaling), but an opposite pattern in human OPCs in AL/CALs.

### LPC and cuprizone demyelination induce transcriptionally similar DAM states that persist through remyelination

In addition to the OL lineage, we also observed dramatic changes in microglia number and state following demyelination (Figure 1d), so we next investigated whether LPC and CPZ elicit different microglia responses. To address this, we first subclustered and reannotated the mouse microglia and macrophages, identifying 5 microglia subclusters (Mg_A-E), PVMs, and a cycling cluster (Figure 5a, Supplementary Figure 10a). Using a previously identified innate immunity gene set, we calculated a gene score and found enrichment of reactive microglia, termed DAMs, within the highly-related Mg_C, D and E clusters, PVMs, and cycling cells (Figure 5a, Supplementary Figure 9b) ^30,31^. Further confirming this activated state, these clusters are enriched for known DAM markers (e.g. *Apoe, Cd63, Lyz2)* and downregulate homeostatic microglia genes (e.g. *Tmem119* and *P2ry12)* (Figure 5b, Supplementary Figure 9c). PVMs express lineage markers *C1qa* and *Trem2,* but lack microglia-specific genes such as *Tmem119* and *Cxcr1*. Notably, PVMs also share expression of many DAM markers, and additionally enrich for immune signaling genes *Igqap1*, *Ms4a7,* and *Msrb1*.

**Figure 5:**
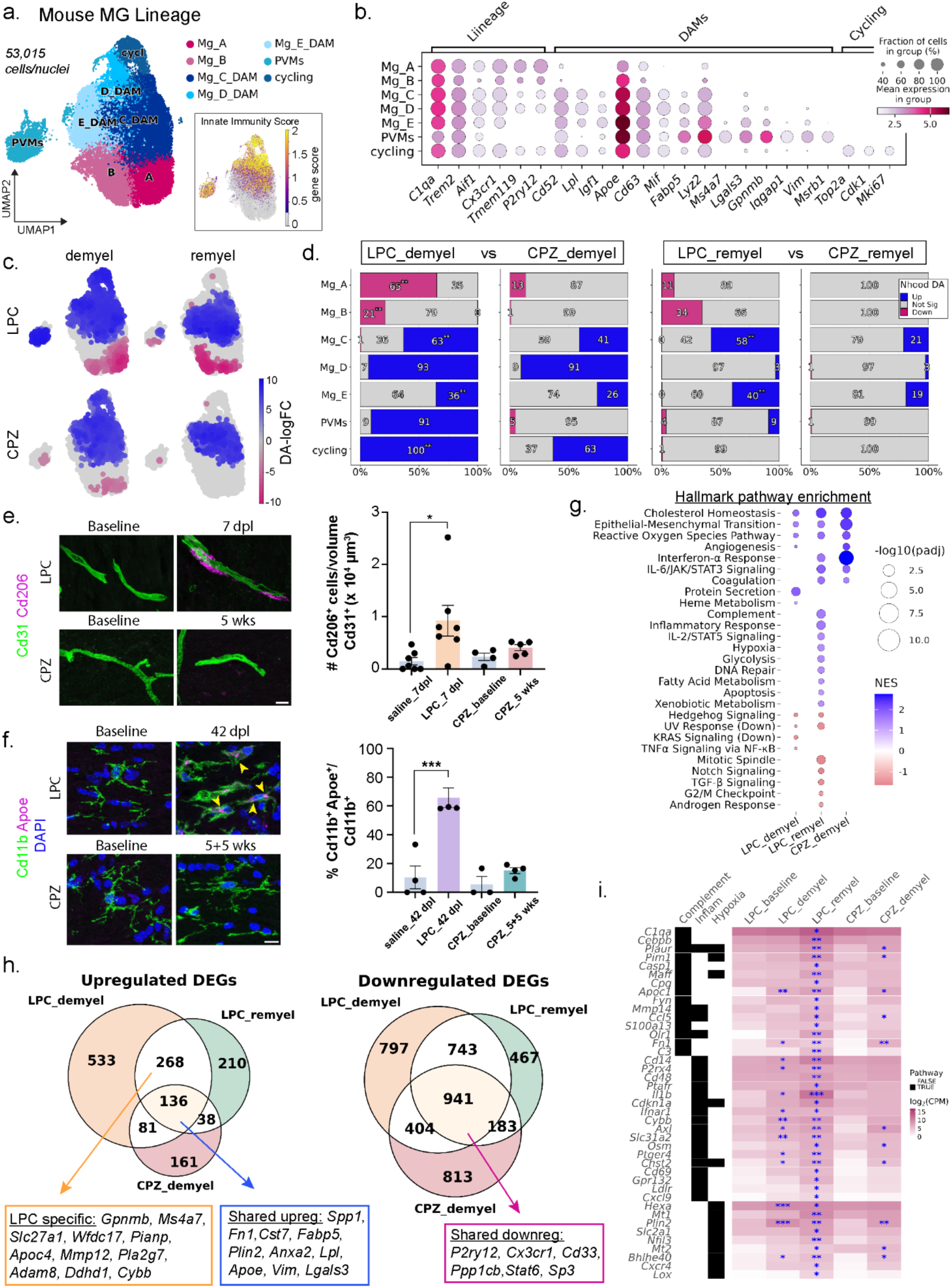
Microglia share conserved gene expression changes in mouse demyelination models, but LPC leads to increased inflammation. a. UMAP embedding of the subset and reclustered mouse macrophage/microglia lineage cells. The inset UMAP shows a gene score for innate immunity-associated genes. (Mg, microglia; DAM, damage-associated microglia). b. Dot plot of lineage- and cluster-specific marker genes across cell types. c. Differential abundance of nhoods superimposed on the UMAP embedding. Each node represents a nhood and is colored by the logFC relative to baseline; non-significant nhoods are shown in grey (spatial FDR < 0.1). d. Bar plot showing the proportion of non-significant (grey), increased (blue), and decreased (magenta) nhoods, as defined in panel c, for each cell type annotated in panel a. Significance annotations (* p < 0.05, ** p < 0.01) indicate results from cell-type-specific enrichment tests comparing LPC vs. CPZ at demyelination (left) and remyelination (right) using Fisher’s exact or chi-squared tests. Exact p-values are provided in Supplementary Table 9. e. Representative IHC images (left) showing Cd31 and Cd206 expression in the corpus callosum. LPC baseline = saline-injected at 7 dpl; CPZ baseline = normal diet. (Scale bar = 10 μm). Bar chart shows the number of Cd206^+^ PVMs per Cd31^+^ vasculature volume. Bars show group means, dots show individual animals, and error bars indicate standard error of the mean (SEM). Statistical significance was assessed by one-way ANOVA with Tukey’s test (\**p* = 0.0229). f. Representative IHC images (left) showing Cd11b and Apoe expression in the corpus callosum. LPC baseline = saline-injected at 42 dpl; CPZ baseline = normal diet. (Scale bar = 10 μm). Bar chart shows the percent of Apoe^+^ microglia (Cd11b^+^Apoe^+^/Cd11b^+^). Bars show group means, dots show individual animals, and error bars indicate standard error of the mean (SEM). Statistical significance was assessed by one-way ANOVA with Tukey’s test (\*\*\**p* = 0.0002). g. Dot plot of Hallmark pathway enrichment across per lesion type. Dot color indicates normalized enrichment score (NES), and size reflects –log10(adjusted *p*). h. Venn diagram showing the unique and overlapping sets of significantly upregulated (logFC ≥ 1.5, padj < 0.05, left) and downregulated (logFC ≤ 1.5, padj < 0.05, right) genes in microglia at the indicated timepoints. Colored boxes highlight DEGs associated with the indicated overlapping gene sets. i. Heatmap showing gene expression changes in LPC remyelination enriched pathways. Heatmaps are colored by the average log₂-transformed CPM expression values. Gene–pathway associations were assigned to pathways using the Hallmark sets and are shown in the left annotation panel, where black tiles indicate pathway membership. Genes are hierarchically ordered first by pathway inclusion and then by average expression across all conditions. Blue asterisks indicate genes with significant upregulation (* for *p* < 0.05, ** for *p* < 0.01, *** for *p* < 0.001).

To compare the distribution of these microglia states between LPC and CPZ demyelination, we performed differential abundance testing using Milo (Supplementary Data 9). LPC demyelination had significantly decreased proportions of homeostatic microglia clusters (Mg_A and B) compared to CPZ demyelination (Figure 5c,d), suggesting stronger microglial activation following LPC injection. Both LPC and CPZ demyelination increased the proportion of DAMs (Mg_C-E) by similar levels, but LPC demyelination induced a significantly higher proportion of cycling microglia (Figure 5d). Interestingly, PVMs were only increased in LPC demyelination, which we confirmed through quantification of Cd206^high^ cells adjacent to blood vessels within the healthy and demyelinated WM (Figure 5e). Both cycling microglia and PVMs were only enriched at demyelination timepoints, suggesting their resolution or clearance promotes remyelination. While DAM states Mg_C and E persisted at remyelination timepoints in both models, they remained significantly higher in LPC compared to cuprizone (Figure 5d). To see if DAMs persist at even later remyelination timepoints, we quantified the percentage of Cd11b^+^ microglia that express the DAM marker, Apoe, at 42dpl for LPC-injected mice and 5 weeks after CPZ removal. Interestingly, we found a high percentage of microglia continued to express Apoe even at late stages of LPC remyelination, whereas microglia no longer expressed Apoe at similar stages of CPZ remyelination (Figure 5f). Similar to our OL analysis, this suggests different timelines of recovery between the two models, with LPC microglia taking longer to return to a homeostatic state.

We next wondered if LPC and CPZ demyelination induce similar changes in gene expression within microglia/macrophages. We tested for DEGs using a pseudobulk approach, comparing LPC and CPZ de- and remyelination microglia to their respective baseline controls (Supplementary Data 10). As PVMs are not present in sufficient numbers in our baseline controls, we compared them with LPC homeostatic microglia. We found large numbers of DEGs across all comparisons, with the largest changes occurring in PVMs and microglia at LPC demyelination (Supplementary Figure 10d). GSEA analysis revealed enrichment of several pathways in common between LPC and CPZ DAMs/PVMs, including cholesterol homeostasis, epithelial-mesenchymal transition, and reactive oxygen species (Figure 5g, Supplementary Figure 10e). In addition, LPC remyelination and CPZ demyelination DAMs were both enriched for immune response pathways (e.g. IFN⍺ response and IL6 signaling) (Figure 5g). We quantified the number of shared DEGs, and found 136 genes were upregulated by DAMs across both models and timepoints, including many well-established DAM genes (e.g. *Apoe*, *Lgals3*, *Spp1*) (Figure 5h). Approximately 67% of upregulated DEGs in CPZ DAMs were also upregulated in LPC DAMs, whereas only 22-27% of upregulated DEGs in LPC DAMs were shared with CPZ, highlighting a LPC-specific DAM response, which includes many immune-related genes (e.g. *Gpnmb*, *Ms4a7*, *Mmp12*). We also found 941 shared downregulated genes across all conditions, which includes homeostatic microglia genes (*P2ry12*, *Cx3cr1*) and inhibitors of phagocytosis (*Cd33*, *Gpr34*), indicating a highly conserved gene downregulation in microglia across both models (Figure 5h, Supplementary Figure 10f). Interestingly, LPC remyelination DAMs were also significantly enriched for inflammatory-related genes (*Il1b, Cxcl9, Cd69*), complement (*C1qa, C3*), and hypoxia/glycolysis (*Mt1*, *Slc2a1*) (Figure 5i). This suggests an interesting transition in DAM states from demyelination to remyelination timepoints, when microglia are important for OPC proliferation, migration, and differentiation^32–34^. The upregulation of complement genes, which are involved in debris clearance, with hypoxia/glycolysis genes suggests a continued pro-inflammatory microglial state at LPC remyelination stages ^13,35–40^.

Overall, LPC induces a stronger microglial response than CPZ, characterized by PVM infiltration, greater microglial proliferation, sustained activation throughout remyelination, and increased expression of inflammatory genes. In contrast, CPZ-induced demyelination elicits a more gradual response over weeks, leading to weaker DAM enrichment and a near-complete return to homeostasis by remyelination. Despite these differences, both models exhibit a conserved DAM state with downregulation of core homeostatic genes.

### Microglia enriched in MS exhibit heterogeneous states across lesion types

To compare the mouse microglial response to demyelination with human microglia in MS tissue, we next characterized microglia states across MS lesions. Reclustering the human microglia revealed 7 subclusters, which we labeled Mg_A-F and PVMs (Figure 6a, Supplementary Figure 11a). Scoring of pro-inflammatory gene sets revealed no enrichment in Mg_A, low enrichment in Mg_E, and strong enrichment in Mg_F, suggesting heterogenous DAM states in MS (Supplementary Figure 11b). Consistently, marker gene expression shows high expression of microglia lineage genes and low levels of inflammatory markers in human Mg_A, which we consider homeostatic microglia (Figure 6b, Supplementary Figure 11c). In contrast, expression of genes involved in activated states are evident in the remaining microglia clusters, including *MYO1E*, *ALCAM*, *APOE*, and *HIF1A*. Human Mg_B and C clusters maintain expression of homeostatic marker *P2RY12* with lower levels of DAM markers, suggesting these clusters represent transitioning microglia. Human Mg_D strongly expresses genes previously used to describe ‘foamy’ microglia, such as lipid storage genes *LPL* and *PPARG* and the lysosome gene *CTSL,* supporting a role for these cells in clearance of myelin debris ^13^. Human Mg_E expresses similar gene expression to Mg_D, but also expresses *NUPR1*, a gene that is critical for repressing ferroptosis ^41,42^. Human Mg_F is more distinct from Mg_C and D, expressing the highest levels of *VEGFA* and *FOSL2*. Finally, the human PVM cluster shows high and specific expression of macrophage markers such as *F13A1*, *MRC1*, *CR1*, and *LYVE1*. Together, these findings suggest that MS induces greater cellular heterogeneity in the activation states than we observed in mouse demyelination models.

**Figure 6:**
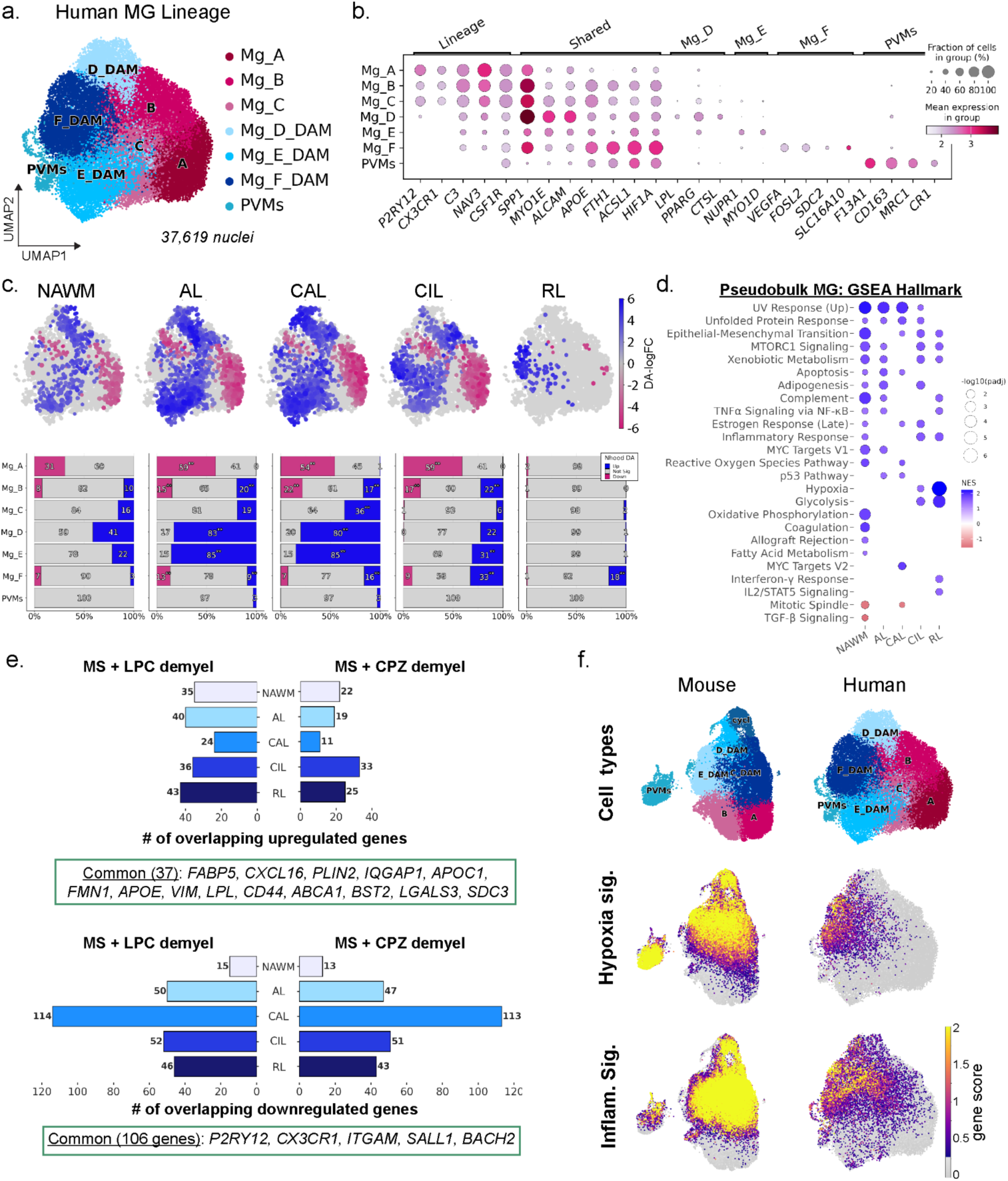
MS lesions enrich distinct microglial states that share core transcriptional features with mice. a. UMAP embedding of the subset and reclustered human microglia/macrophage lineage cells. b. Dot plot of lineage- and cluster-specific marker genes across cell types. c. UMAP plots showing differential abundance nhoods for each lesion type relative to WM, computed using Milo. Blue indicates significantly increased nhoods, magenta indicates significantly decreased, and grey indicates non-significant (spatial FDR < 0.1). Bar plots (bottom) showing the proportion of non-significant (grey), increased (blue), and decreased (magenta) nhoods for each cell type. Significance annotations (* for *p* < 0.05 and ** for *p* < 0.01) indicate results from a separate statistical test assessing cell-type-specific enrichment using pairwise Fisher’s exact or chi-squared tests comparing the lesions to NAWM. Exact p-values are provided in Supplementary Table 11. d. Dot plot of Hallmark pathway enrichment across per lesion type. Dot color indicates normalized enrichment score (NES), and size reflects –log10(adjusted *p*). e. Bar charts showing overlaps of upregulated (top) and downregulated (bottom) MG DEGs in LPC (left) and CPZ (right) at demyelination compared with MS lesion types. The green boxes indicated common genes that are up or downregulated in both LPC and CPZ datasets, and at least 1 of the MS lesion conditions. f. UMAP plots of mouse (left) and human (right) MG cells and gene scores derived from genes involved in hypoxia and inflammatory response that are upregulated in LPC remyelination and at least 1 MS lesion type.

We next investigated whether MS lesions differentially enrich for activated microglia states, using Milo analysis and statistical testing (Supplementary Data 11). While shifts from homeostasis were clearly evident in the NAWM microglia, consistent with previous reports showing reactive microglia throughout the CNS in MS patients, the changes in cell proportions were far fewer compared to the lesion conditions (Figure 6c) ^41^. MS lesions had significantly decreased proportions of homeostatic microglia (Mg_A) across lesion types, except for RLs (Figure 6c). ALs and CALs were strongly enriched for DAM clusters Mg_D and Mg_E, suggesting foamy microglia are induced in regions of high myelin debris. In contrast, CILs and RLs were enriched strongly for DAM cluster Mg_F, with reduced enrichment of DAM clusters Mg_D and E. This raises the possibility that remyelination-specific microglia may create an environment conducive to repair. Mg_F expresses *VEGFA*, which could promote angiogenesis in MS, and has been shown in mouse models to enhance OPC survival and differentiation ^43,44^. We observed weak enrichment of the PVM cluster, which comprised very few cells in the analysis, in ALs and CALs.

To define the transcriptional differences in activated human microglia across distinct lesions, we performed pseudobulked microglial differential gene expression testing, comparing all MS lesions to control WM (Supplementary Data 12). We found several hundred genes differentially expressed in each lesion type, with NAWM microglia containing the smallest changes (369 total DEGs) and CAL microglia containing the highest (955 total DEGs) (Supplemental Figure 12a). Among them, only 27 upregulated and 20 downregulated genes were shared across all lesion types, but microglia in ALs and CALs shared an additional 32 upregulated and 62 downregulated genes, whereas CILs and RLs shared 61 upregulated and 17 downregulated genes (Supplementary Figure 12b). GSEA pathway analysis revealed that several pathways were upregulated in microglia across multiple lesion types, including UPR (*BAG3, DNAJA4, TSPYL2*), MTORC1 signaling (*HSPD1*, *HSPA4*, *SERPINH1*), and epithelial-mesenchymal transition (*TGFBI, FERMT2, FLNA*) (Figure 6d). Interestingly, CIL and RL microglia were also enriched for glycolysis (*PKM, PYGL*) and hypoxia-related genes (*VEGFA*, *PLIN2*, *FOSL2*, *NDRG1*), which mirrors mouse microglia in LPC remyelination samples.

### Mouse demyelination models recapitulate core microglia activation states observed in MS lesions

We then compared microglia from LPC and CPZ demyelination models with microglia across the MS lesion types. MetaNeighbor analysis showed that cell types from both species grouped into two main clusters, representing homeostatic microglia and DAMs/PVMs (Supplementary Figure 12c). Within the DAMs/PVMs branch, we identified two subclusters, one containing mouse and human Mg_E,F and PVMs, and one containing mouse and human Mg_D and cycling microglia (Supplementary Figure 12c). We found that mouse DAMs share some overlap of upregulated genes with the human DAMs, with genes such as *APOE, VIM, LPL,* and *FABP5* upregulated by microglia in MS lesions, LPC, and CPZ demyelination (Figure 6e). This reveals a common core of genes that are highly conserved in activated microglia across species. LPC de- and remyelination DAMs shared slightly more upregulated genes with MS microglia (24-43 and 26-42 DEGs across lesion types, respectively) than CPZ demyelination DAMs (11-33 DEGs across lesion types). Both LPC and CPZ DAMs shared similar numbers of downregulated genes with the human DAMs across MS lesions, though notably, CAL microglia shared the highest number of downregulated genes with mouse DAMs (Figure 6e, Supplementary Figure 12d).

Overlap coefficients confirmed strong within-species similarity, particularly among MS lesion comparisons (AL/CAL and CIL/RL), with high Spearman correlations (>0.716 in human and >0.601 in mouse), indicating well-conserved logFC direction and magnitude. Across both mouse models, downregulated gene overlap with MS DAMs remained consistent (0.34–0.45), while upregulated gene overlap was higher in LPC de- and remyelination than in CPZ demyelination. Spearman correlations between mouse models and MS DAMs were moderate overall (0.22–0.29), though comparisons to the CAL state were uniformly lower (0.05–0.13). Together, these analyses indicate that both demyelination models share a modest but reproducible transcriptional similarity with MS microglia, particularly in the downregulation of homeostatic programs, with LPC models more closely mirroring MS-associated inflammatory activation.

To further explore the similarity between LPC remyelination DAMs and RL DAMs, we next calculated gene signature scores based on upregulated hypoxia- and inflammation-associated gene sets. Both the hypoxia and inflammation gene signatures were most strongly enriched in the human Mg_F DAM state that is present in RLs (Figure 6f). In contrast, both gene signatures were broadly upregulated across the mouse DAM populations, rather than restricted to a specific subcluster. Therefore, we conclude that mouse demyelination models capture core conserved features of microglial activation seen in MS, including loss of homeostatic identity and induction of a shared inflammatory program. However, the overall DEG overlap and gene signature patterns support that human MS microglia engage additional activation pathways and cell-state diversity not found in mouse models. This may reflect the differences between acute demyelination mouse models versus chronic human disease. Alternatively, this may suggest that mouse microglia perform multiple functions required to clear myelin debris and create a permissive environment for remyelination, but these functions may be diversified in humans and restricted to specific subtypes, consistent with previous reports of evolutionary divergence in human microglia ^45^.

## Discussion

The use of animal models to study demyelination and remyelination has been critical in advancing our understanding of OL susceptibility in neurodegenerative disease, and has been a valuable tool to address the urgent, unmet need for therapies promoting myelin protection and repair. Toxicity models, including CPZ and LPC, offer precise temporal control over damage and a predictable timeline for repair, making them particularly well suited for assessing therapies for halting disease progression and promoting remyelination. Numerous preclinical therapies have been identified using these models, with a 2019 report stating that ∼23% have entered clinical trials and 5 received approval (Fingolimod, Fumaric acid ester, Glatiramer acetate, Methotrexate, Methylprednisolone)^46^. However, whether distinct cellular responses are induced by different demyelination treatments remains unexplored, and these models have not been systematically compared to human patient samples across multiple cell types. Therefore, we analyzed single-cell/nuclei datasets to profile transcriptional changes in LPC- and CPZ-induced demyelination and remyelination, comparing them to the largest MS dataset to date.

We found that CPZ demyelination induces a distinct DAO phenotype that is not observed at significant numbers in LPC demyelination. This DAO state is enriched for cellular stress pathways, including UPR and DNA damage, with significant upregulation of *Cdkn1a* and *Nupr1* genes. *Cdkn1a* (p21) expression is frequently induced following DNA damage and its upregulation is linked to cellular senescence, suggesting that CPZ may push OLs towards a senescent cell state^47^. While the function of *Nupr1* in oligodendrocytes has not been examined, it is known to protect against ferroptosis in cancer cells ^42,48,49^. Markers of ferroptosis have been found upregulated in MS lesions, mostly within microglia ^50–54^, but our results suggest a role for Nupr1 within oligodendrocytes as well. Interestingly, Nupr1 has also been linked with cellular senescence in MS lesions ^55^. Importantly, the CPZ DAO state exhibits greater DEG overlap with DAOs in the EAE model and MS lesions than LPC-induced states. Many pathways that are increased in CPZ DAOs (UPR, Myc targets, mTORC1) are also upregulated in human DAOs within RLs, which could reflect the ongoing de/remyelination within the mouse model and the different cell-cell signaling that may occur across the different MS lesion types. As OPCs and DAMs in the mouse toxicity models also share similar signaling pathways with human OPCs or DAMs within CILs/RLs, it suggests that CPZ and LPC recapitulate the remyelinated MS white matter environment, rather than active lesions.

Despite the unique CPZ demyelination DAO state, both mouse models converge onto a similar remyelination state that continues to express DAO genes (*Serpina3n, C4b*), suggesting that newly-generated mouse OLs exist within a stressed state during remyelination. In addition, we found that this remyelination DAO state expresses immune-related genes, including *Socs3*, *H2-D1*, and *B2m*. While the function of these genes in OLs remains to be determined, we feel this has several important implications – first, upregulation of MHC genes in OL lineage cells is not unique to the EAE model, suggesting that non-T cells may induce this response. IFN signaling has been shown to induce expression of MHC molecules in OPC cultures ^27^, so we hypothesize that secretion of IFN by microglia or other cell types in the LPC and CPZ models induces this response. Second, the functional consequences of this MOL state at remyelination remain unclear. For example, it is not known if these OLs produce functional myelin that matures over time. A recent study demonstrated that even incomplete remyelination following CPZ-induced demyelination is sufficient to restore neuronal function^2^. However, whether restoring proper MOL heterogeneity is essential for functional tissue regeneration remains an open question, requiring further investigation into how transcriptional phenotypes translate to MOL structure and function. Finally, the maintenance of *Socs3* expression at late remyelination timepoints in LPC, but not CPZ, mirrors the sustained DAM activation at LPC remyelination timepoints. Whether this prolonged inflammatory response is beneficial for remyelination remains unclear, but microglia have been previously shown to promote remyelination independent of debris clearance^32–34^. As Socs3 is a suppressor of cytokine signaling, inhibiting NF-κB and JAK/STAT signaling, its upregulation may be a compensatory mechanism by OLs in response to the inflammatory environment. Interestingly, Socs3 knockout mice displayed reduced demyelination following CPZ feeding in a previous study, suggesting that inhibiting Socs3 may promote OL survival ^56^. As OL survival has been linked as a critical step of remyelination ^57^, determining if ablation of Socs3 after demyelination increases OL survival is an important next step, especially as we found upregulation of *SOCS3* in human OLs in CILs.

A benefit of the human MS dataset that we use in our analysis is that it contains samples across different lesion types, whereas other studies largely focus on CALs. While the original publication of this human dataset presents cellular state changes across these different lesion types, it was not the primary focus of their study ^14^. Interestingly, we find that DAO states do not change across the MS lesion types, but both OPCs and DAMs exhibit wide heterogeneity. OPCs, in particular, show an interesting inverse pattern in gene signaling pathways between AL/CALs and CIL/RLs. OPCs in active MS lesions upregulate inflammatory genes, which may play a role in differentiation arrest and promote chronic inflammation, suggesting strategies to ameliorate these detrimental changes may be clinically relevant ^11,15,28^. However, our data suggests that there is a diversity of OPC states in addition to immune expression that need to be further studied in MS lesions to provide effective targets for therapies. Importantly, mouse and human OPCs have the least number of overlapping genes across the different cell populations we explored, indicating that mouse models do not recapitulate human OPC pathology in MS very well. One potential explanation is the more advanced age of human OPCs in post-mortem tissue, compared to young adult mice that are commonly used for experiments. As mouse OPCs undergo transcriptional and functional changes with aging ^58^, it would be interesting to compare OPCs following demyelination in older mice.

Microglia and macrophages in the MS cohort also show important differences from the mouse models, particularly in their increased heterogeneity. Unlike in mouse models, human microglia do not show evidence of proliferation, aligning with previous findings that microglial populations are largely stable, with an estimated lifespan of ∼4 years ^59^. We identify a transcriptional signature similar to previously described foamy microglia (Mg_D), which are highly enriched in AL and CAL lesions. In contrast, microglia in remyelinating regions are largely homogeneous for the Mg_F state, which is characterized by strong enrichment for hypoxia and inflammatory signaling. While some differentially expressed genes and pathways are conserved between human and mouse microglia, there is no clear enrichment for a specific human microglial subtype in the mouse models. Instead, the shared response primarily involves canonical demyelination-activated genes, such as *LPL, GPNMB,* and *APOE*. One potential explanation for the more homogeneous microglial response in mice is the shorter remyelination timeline (∼3 weeks), compared to the prolonged disease course in MS. In longer-term neurodegenerative models, such as AD, distinct temporal microglial activation states emerge^60^. Comparing microglial responses in chronic or repeated demyelination models could reveal heterogeneity not captured in short-term studies and better align with the diversity observed in MS.

Altogether, our study suggests that LPC and CPZ models should be used strategically to investigate pathways enriched in each respective condition. We propose that studies focused on OL dysfunction, pathology, and/or survival in MS use the CPZ model, for it more closely recapitulates the human DAO state. However, OLs in both models do share some signaling pathways, such as TNFɑ signaling, as well as a similar remyelination state. TNFɑ signaling is also increased in MS lesions and blocking soluble TNF improved outcomes in EAE, suggesting that elevated TNFɑ signaling may have detrimental effects in both LPC and CPZ models ^61^. However, blocking TNFɑ signaling has been found to increase the risk of MS ^62^, as well as worsen disease ^63^, highlighting the complexity of this pathway in MS disease progression. As both LPC and CPZ induce strong microglial activation phenotypes, with overlapping transcriptional changes affecting hundreds of genes, our results support that both models are suitable for assessing microglial dynamics in response to demyelination. However, LPC induces the strongest increase in cycling microglia and sustains microglial activation longer during remyelination, thereby providing a more robust model for testing potential therapies for reducing microglia activation. While no current MS therapies directly target microglia, some drugs with off-target microglial effects show regenerative benefits, suggesting that targeting microglia directly could enhance remyelination^59^. LPC demyelination also induces a PVM response that we do not observe in CPZ, so it may also serve as the preferred model for distinguishing microglia versus macrophage function across the span of injury and repair. While it remains difficult to definitively identify CNS macrophages versus microglia in post-mortem MS tissue, previous work has found evidence of increased macrophage infiltration in active MS lesions, where they are hypothesized to contribute to chronic inflammation^64^. One limitation of our study was insufficient astrocyte numbers for robust downstream analysis, as astrocytes are known to play a critical role in both demyelination models^17,18^. Future studies comparing astrocyte states between mouse models and MS lesions are necessary to fully understand the signaling mechanisms driving remyelination. For example, astrocytes have been shown to coordinate debris clearance via crosstalk with microglia^61^. Future studies investigating how cell-cell interactions regulate remyelination at the tissue level will be essential for understanding the resolution of demyelination and the mechanisms driving effective myelin repair.

## Closing information

### Author information

E.L.A. performed the analysis of the single cell/nuclei RNA-seq datasets ,and generated the figures. V.R.V., E.A.M. and M.M.H. performed the RNAscope/immunohistochemistry (IHC) validation experiments. K.L.A. performed the single nuclei RNA-seq experiments and supervised the project. E.L.A. and K.L.A. conceptualized the project and wrote the manuscript.

## Acknowledgements

We would like to thank Dr. Vittorio Gallo at Seattle Children’s Hospital for his mentorship of K.L.A. during early stages of this project and Dr. Cody Smith at the University of Notre Dame for critically reading this manuscript. We thank Dr. Bharat Mishra for analysis and statistical advice. We thank the Freimann Life Science Center at the University of Notre Dame for mouse breeding and husbandry, and Dr. Sara Cole from the Notre Dame Integrated Imaging Facility for her microscopy support. Images presented in this manuscript were generated using the instruments at the NDIIF/Optical Microscopy Core. We thank Karuna Panchapakesan, Dr. Susan Knoblach, and Dr. Surajit Bhattacharya at Children’s National Hospital for their assistance with single nuclei RNA-seq library generation and analysis. Finally, we thank members of the Adams lab for their discussion. This work was supported by a career transition award to K.L.A. from the National Multiple Sclerosis Society.

## Competing interests

The authors declare no competing interests.

## Data availability

All sequencing data generated in association with this study are available in the Gene Expression Omnibus (GEO) under accession number GSE293850. The additional data used in this study includes GSE182846, and GSE148676. The human MS dataset is available on Zenodo (DOI: 10.5281/zenodo.8338963).

## Code availability

Details of the analyses used herein are described in the Methods sections below, and jupyter notebooks are available at https://github.com/AdamsLabNotreDame. Any additional information required to reanalyze the data reported in this paper is available from the lead contact upon request.

## Methods

### Experimental procedures

#### Animals

Wild-type C57BL/6 mice were maintained in the animal facility at the Children’s National Hospital and the University of Notre Dame according to the Institutional Animal Care and Use Committee and the National Institutes of Health guidelines. Both male and female mice were analyzed for all experiments. Mice were maintained in the animal facility under a 12 h dark–light cycle, and constant temperature (20–26 °C) and humidity maintenance (40–60%). All studies complied with all relevant animal use guidelines and ethical regulations.

#### Demyelination protocols

For LPC demyelination, adult (8-12wk) C57BL6 mice were anesthetized with ketamine/xylazine, the skull exposed, and a small hole drilled 1 mm rostral and 1 mm lateral of bregma. The animal was placed in a stereotactic rig and injected with 1uL of 1% LPC into the corpus callosum of one brain hemisphere at 2 mm depth over 5 minutes. Saline was injected on the contralateral side using the same steps as above. Mice were sacrificed as described below at 7 dpl for snRNA-seq and RNAscope/IHC for the demyelination timepoint, at 28 dpl for RNAscope/IHC at the remyelination timepoint, and at 42 dpl for the late recovery timepoint.

For snRNA-seq, adult (8-12wk) CPZ demyelination was conducted by feeding adult (8-12wk) C57BL6 mice 0.2% CPZ pellets (Envigo TD.140803) for 5 weeks. For RNAscope/IHC validation, mice were fed 0.3% CPZ pellets (Envigo TD.140805) for 5 consecutive weeks, and then either sacrificed as described below at the end of 5 weeks for the demyelination timepoints, or returned to normal chow and collected 3 or 5 weeks later for the remyelination and late recovery timepoints. During the study, the lab switched from 0.2% to 0.3% CPZ pellets to improve the efficiency of demyelination. Control mice were fed normal chow and processed as described below.

#### Dissection and single nuclei preparation

For LPC demyelination snRNA-seq collection at 7 dpl, mice were IP injected with neural red dye 2-3 hrs prior to tissue collection to label the lesion within the corpus callosum. Mice were then anesthetized, perfused with cold PBS for 5 mins, and the brain was dissected out into cold PBS. Coronal slices of the brain were made, to identify the lesion location, and the corpus callosum lesions were dissected out. For LPC controls, the corresponding contralateral side with saline injection was also collected. For CPZ fed mice, the whole corpus callosum was dissected out. Tissue was placed in Eppendorf tubes, excess liquid was removed as much as possible, and tissue was flash frozen in liquid nitrogen. Frozen tissue was stored at −70°C.

Nuclei were pooled from 5 female and 5 male mice per condition for each replicate, with two replicates per condition, (LPC: baseline = saline injected, demyelination = 7dpl, CPZ: baseline = normal chow, demyelination = 5 wks of 0.2% CPZ). Pooled tissue was transferred to chilled glass dounce with 1mL of lysis buffer + 1% TritonX100 + 1 μM DTT (Sigma Nuclei Pure Prep Kit, Cat #NUC-201). Samples were dounced on ice 10 times with pestle A, then 10 times with pestle B. The suspension was passed through a 40μM FlowMi cell strainer. Dounces were rinsed with 1mL of low sucrose buffer and passed through the cell strainer as well. Samples were centrifuged at 300 rcf, at 4°C, for 5 mins in a fixed-angle centrifuge. Supernatant was carefully decanted, and the pelleted nuclei were washed with 1.5mL of low sucrose buffer, and centrifuged again at 300 rcf, at 4°C, for 5 mins. Pelleted nuclei were then resuspended in 500μL Nuclei wash buffer (PBS + 1% UltraPure BSA + 0.2U/μL Protector RNase inhibitors) + 900μL of 1.8M sucrose cushion buffer, then carefully layered on top of 500μL 1.8M sucrose cushion buffer in a 2mL low-bind tube.

Samples were centrifuged at 13,000 rcf, at 4°C, for 45 mins. Supernatant was removed, and nuclei were resuspended in 1mL of Nuclei wash buffer, passed through a 40μM FlowMi cell strainer and counted using a Cell Countess II. Approximately 6,000 nuclei were loaded per sample onto the 10X chromium controller per manufacturer’s instructions. Single nuclei libraries were generated using the 10X single cell 3’ reagents and protocol (v3.1). Libraries were sequenced on an Illumina Novaseq6000, with an average number of 80,000 reads per cell for each sample.

#### EdU treatment and detection

LPC-injected or CPZ-fed animals received intraperitoneal injection of EdU (Sigma, 900584-50MG) at 50mg/kg. For LPC animals we injected EdU at 6, 9 and 12 dpl, and for CPZ animals we injected Edu at 2, 5 and 8 days after returning them to a normal diet. The RNAscope Multiplex Fluorescent Reagent Kit V2 (Advanced Cell Diagnostics #323100) was used to perform fluorescent labeling of the indicated RNA molecules according to the kit manual. RNAscope probes used were: *Olig2* (447091-C3) and *Socs3* (437741). Following RNAscope, EdU was detected using the Click-iT EdU Cell Proliferation Kit (Thermo, C10637) following the manufacturers instructions. Sections were then coverslipped with mounting media (Southern Biotech). Sections were imaged using the Leica Stellaris 8 DIVE confocal microscope and cell counting was analyzed for at least 3 sections per animal. Cells with at least two fluorescent puncta were counted as positive for that probe.

#### RNAscope and IHC of OL lineage

Mice were anesthetized using isoflurane and intracardially perfused with cold PBS followed by 4% PFA for 5 minutes each. The brain was dissected out into cold 4% PFA and postfixed overnight. The tissue was then cryoprotected in 30% sucrose in PBS. Brains were embedded in OCT and frozen on dry ice. Sections were collected onto slides (12µM) and stored at −70°C, or as floating sections (40µM) and stored in PBS at 4°C.

IHC was performed first by thawing the slides (all steps are performed at RT unless otherwise indicated) in a humidifying tray for 5mins, permeabilized for 15 mins with 1xPBS + 0.1% tween (PBST), and blocked in blocking buffer (PBST + 1% normal donkey serum (NDS)) for 30 mins. Primary antibody (rabbit anti-Olig2, Millipore AB9610, 1:300 or goat anti-Olig2, R&D Systems AF2418, 1:100) was incubated in blocking buffer overnight at 4°C. Slides were washed 3 times for 5 mins each with PBST, and incubated with the indicated fluorophore-conjugated secondary antibodies (AlexaFluor) in blocking buffer for 2 hrs. Tissue was washed again 3 times with PBST, and then post-fixed with 4% PFA for 10 mins. Next, the RNAscope Multiplex Fluorescent Reagent Kit V2 (Advanced Cell Diagnostics #323100) was used to perform fluorescent labeling of the indicated RNA molecules according to the kit manual, beginning with protease treatment. RNAscope probes used were: *Cdkn1a* (408551-C2), *Socs3* (437741), *Pdgfra* (480661-C3), *Nupr1* (434811), and *B2m* (415191-C2). Following RNAscope, sections were coverslipped with mounting media (Southern Biotech). Sections were imaged using the Leica Stellaris 8 DIVE confocal microscope and cell counting was analyzed for at least 3 sections per animal. Cells with at least two fluorescent puncta were counted as positive for that probe. Olig2^+^ cells positive for *Pdgfra* were counted as OPCs, and Olig2^+^ cells negative for *Pdgfra* were counted as MOLs.

#### IHC analysis of microglia/macrophages

Mouse brain tissue was collected as described above. For analysis of PVMs, 40µm floating sections were permeabilized for 15 mins with 1xPBS + 0.1% tween (PBST), and blocked in blocking buffer (PBST + 5% NDS) for 1 hr. Primary antibodies (goat anti-CD206, R&D Systems AF2535, 1:400 and rat anti-Cd31, BD Pharmingen 550274, 1:100) were incubated in blocking buffer overnight at 4°C. Tissue sections were washed 3 times for 5 mins each with PBST, and incubated with the indicated fluorophore-conjugated secondary antibodies (AlexaFluor) in blocking buffer for 2 hrs. Sections were washed again 3 times with PBST, and then mounted on slides with Southern Biotech mounting media. For analysis of DAMs, 12µm brain sections on slides were permeabilized for 15 mins with 1xPBS + 0.1% tween (PBST), and blocked in blocking buffer (PBST + 1% NDS) for 30 mins. Primary antibodies (rabbit anti-Apoe, Cell Signaling 49285T, 1:100 and rat anti-Cd11b, Biorad MCA74G, 1:250) were incubated in blocking buffer overnight at 4°C. Tissue sections were washed 3 times for 5 mins each with PBST, and incubated with the indicated fluorophore-conjugated secondary antibodies (AlexaFluor) in blocking buffer for 1 hr. Sections were washed again 3 times with PBST, and then coverslipped with Southern Biotech mounting media. Sections were imaged using the Leica Stellaris 8 DIVE confocal microscope and cell counting was analyzed for at least 3 sections per animal. For analysis of PVMs: The total number of CD206^high^ cells adjacent to CD31^+^ blood vessels within the demyelinated WM was counted and normalized to the volume of CD31 staining, measured by use of surface tool in IMARIS software. For analysis of DAMs: the number of Apoe^+^ Cd11b^+^ cells were counted using FIJI and graphed as a percentage of total Cd11b^+^ cells.

## Data analysis

### Single cell and nuclei raw data processing

<u>Mouse:</u> Demultiplexed fastq files were aligned to mm10 using Cell Ranger (v6.0.1), and published datasets were downloaded from GEO, and mapped using Cell Ranger (v7.2.0) ^8,9^. Mapped h5 files were run through CellBender to remove ambient RNA ^65^. Corrected count matrices for each sample were processed using scanpy tools and recommended best practices by retaining cells for downstream analysis ^66^. Doublet detection and removal was performed using Solo, by first training a scVI model on each batch and then removing predicted doublets ^67^. After doublet removal, data was merged across its publication of origin (ie. unpublished single nuclei data, Pandey et al, or Shen et al). Each of these 3 datasets was then subset based on quality metrics for that dataset- cells or nuclei were removed if they had less than 200 genes identified, more than 20% ribosomal content, or more than 5% mitochondrial reads. Additionally, we filtered the data to remove the highest outliers using the following criteria: the single nuclei data was filtered to remove samples with more than 15,000 total reads, or more than 7,000 genes. Pandey et al single cell data was subset to remove cells with more than 8,000 genes per cell, or 50,000 reads per cell. Shen et al was subset to remove cells with more than 6,000 genes identified, or more than 20,000 reads per cell.

<u>Human:</u> The MS data quality control counts matrix was downloaded from Zenodo (DOI: 10.5281/zenodo.8338963), subset to only the white matter samples, and converted to an h5ad file ^68^. A major advantage of this dataset is that it contains carefully staged lesions. In brief, the subset data comprises 92 samples collected from 47 individuals, including 13 control (Ctr) and 34 MS patients, categorized by disease progression as relapse remittent (RR), secondary progressive (SP), or primary progressive (PP) MS (Supplementary Figure 2D). The WM only subset contains 13 controls (WM, mean age at death: 64.8 years, median: 70 years), 17 normal-appearing white matter samples (NAWM, mean: 61.3 years, median: 66 years), 21 active lesions (AL, mean: 51.5 years, median: 47 years), 17 chronic active lesions (CAL, mean: 54.7 years, median: 48 years), 13 chronic inactive lesions (CIL, mean: 65 years, median: 69 years), and 11 remyelinated lesions (RL, mean: 66.2 years, median: 69 years). Samples were further controlled by removing nuclei with greater than 10% mitochondrial counts, more than 10,000 genes per cell, and more than 50,000 total reads.

### Dataset integration and annotation

<u>Mouse:</u> First, each dataset was integrated within its own experimental group (i.e. each data source separately) to obtain harmonized cell type labels using scVI ^69^. Briefly, we subset the data to the top 5,000 highly variable genes (HVGs), with the batch key set to each individual batch, and the percent counts for mitochondria and ribosome as continuous covariates. The latent representation from the trained scVI model was used to calculate the neighborhood graph, and then the UMAP embedding. Cell types were labeled by visual inspection of known marker gene expression, using markers calculated using scanpy’s rank_genes_groups, and using custom cell type scoring using the marker gene sets detailed in Koupourtidou et al (see also lists under Gene Set Scoring). For full integration of the mouse single cell and nuclei data we used sysVI, which was developed to handle challenging integration tasks, such as between single cell and single nuclei data or between species ^70^. For the mouse only integration we used the top 3000 HVGs, using the scanpy highly_variable_genes function set to ‘cell_ranger’, batch key set to the system variable (single cell or single nuclei), and each of the individual samples as a categorical covariate. We verified marker gene expression was enriched in the cell type labels, and that each cell type contained single cells and nuclei.

<u>Human:</u> To match our mouse corpus callosum datasets, we integrated the WM samples only using scVI, and produced a similar embedding and annotation as the published dataset ^71^. For the mode, the batch key was set to the sample_id (representing each individual sample), and RNA count and percent mitochondrial count were given as categorical covariates. For training the model we used n_layers = 2, n_latent = 30, and gene_likelihood = ‘nb’.

### Differential abundance analysis

Compositional changes in the total UMAP was quantified using scCODA ^20^. For the mouse integration, the samples were split into LPC or CPZ treatment groups and the respective baseline samples used as the control. Pericytes were used as the reference cell type. For the human data, the WM sample was used as the control, and the cluster “Endo + Peri” was used as the reference cell type. The default FDR of 0.05 was used.

Milopy, the python implementation of the Milo toolset, was used to calculate the differential abundance (DA) between neighboring cells in the cell type-specific embeddings ^24^. For the mouse OL lineage, neighborhoods (nhoods) were recalculated using the X_sysVI representation and n_neighbors set to 100, and for mouse microglia set to 300, for human oligodendrocytes set to 200, and for human microglia to 100. The make_nhoods function was applied with proportion set to 10% to create nhoods, and the median was inspected to ensure that there are on average at least 5 cells per sample per nhood. Pairwise comparisons were performed to calculate DA, and build an nhood_graph for each treatment condition compared to the relevant baseline for mouse samples or WM for human MS lesion samples. The nhoods were annotated by the cell type labels by calculating the fraction of cells in a hood, if it was more than 60% a single cell type. For nhoods that were less than 60% a single cell type, the annotation was set to ‘Mixed’. The final calculation and annotation was saved from the milo object, and used for statistical analysis.

To compare the distribution of nhoods across conditions, we aggregated counts of upregulated, downregulated, and non-significant nhoods by cell type and condition. For each cell type and significance category, we constructed contingency tables of observed counts and totals, and applied either Fisher’s Exact Test (for tables with <100 total counts) or Pearson’s Chi-squared Test (for larger tables). Resulting p-values were corrected for multiple testing using the Benjamini–Hochberg procedure. For visualization, we generated stacked bar plots showing the proportions of upregulated, downregulated, and non-significant neighborhoods per cell type in each condition. Significant differences between conditions were denoted with asterisks on the bar with higher proportions of significant nhoods that corresponded to adjusted p-value thresholds.

### Differential expression analysis

<u>Mouse:</u> Differential gene expression was calculated using a pseudobulking approach, summing raw gene expression counts in each replicate for each broad cell type with at least 30 cells. Pseudobulk counts were then subset to test individual pairwise comparisons. Each test was performed separately because not all samples contain single cell and nuclei samples, so batch correction was performed differently. For each pairwise comparison, count matrices were trimmed using the edgeR filterByExpr function with the design matrix as input, and keep.lib.sizes set to FALSE. Normalization factors were calculated using the TMM method. For mixed samples (with cells and nuclei) we used voom with quality weights and duplicate correlation twice, blocked on

the library type ^72^. For samples with only single cells, we used voom with quality weights and no blocking. Using lmFit, a linear model was fitted that included blocking on the batch key and correlation set to the duplicateCorrelation output. Empirical Bayes smoothing was applied, and the results table extracted using topTable with n=Inf to include all genes in the output. The full differential expression output for each cell type tested is included as a supplementary table. Differential testing in the CPZ remyelination timepoint produced poor results due to low cell number (2,407 across 3 replicates), small library sizes, and high heterogeneity between replicates. Unless otherwise noted in the figure legend, genes are considered significantly differentially expressed if the adjusted *p*-value is less than or equal to 0.05, and the absolute logFC is greater than or equal to a 1.5 fold change (corresponding to |logFC| ≥ 0.585). Venn diagrams were made using the matplotlib library venn, and upset plots were made using UpSetPlot in python ^73^. Ggplot was used to create heatmaps of the normalized and transformed counts per million (log₂[CPM+1]) that were averaged across each replicate within a treatment group. Heatmap gene annotations were assigned based on inclusion in the mouse orthology-mapped GSEA Hallmark gene sets.

<u>Human:</u> Pseudobulked samples were prepared as above for the mouse data. Briefly, expression counts in each replicate and each broad cell type with at least 15 cells were pseudobulked, then individual comparisons were subset for downstream analysis. First we removed outlier samples with a log library size 3X median absolute deviation (MAD) below the median, as library size seemed to be one of the strongest drivers of variability. Next, for each comparison, the counts matrices were trimmed using the edgeR filterByExpr function and a design matrix with donor age (as categories of under 50, 50-70 and above 70), sex, and post mortem interval (PMI, as categories of less than 6h, 6-12h or more than 12h), and keep.lib.sizes set to FALSE. The TMM method was used to calculate normalization factors. Differential expression was assessed using generalized linear mixed models (GLMMs) implemented in the glmmTMB R package, assuming a negative binomial distribution (nbinom2) to account for overdispersion. For genes with zero counts in all samples (very few), a count of 1 was added to the sample with the largest library size to enable model fitting. Each model included group (e.g., white matter vs. lesion), age_cat, sex, and PMI as fixed effects, and a random intercept per donor to capture inter-individual variability. For each gene, the model output included the estimated log-fold change for the group of interest, standard error, p-value, convergence status, positive-definite Hessian flag (pdHess), log-likelihood, and AIC. Genes for which the model failed or the Group coefficient was not estimable were flagged as errors. P-values were adjusted for multiple testing using the Benjamini–Hochberg false discovery rate (FDR) procedure. A post hoc quality flag was created (status = “OK” for models with convergence = 0 and pdHess = TRUE), and downstream analyses were restricted to these high-confidence models. The final output is a ranked list of genes with robust evidence of differential expression between conditions.

### Gene set enrichment analysis

Fast gene set enrichment analysis (fgsea) was implemented to score pathway enrichment ^74^. For the mouse datasets, the ranking score was set to the sign of the logFC multiplied by −log10(*p*), assigning positive values to upregulated genes and negative values to downregulated genes, with larger magnitudes reflecting greater statistical significance. For the human dataset, *p*-values of exactly zero were replaced with a minimal value to prevent numerical errors. Then, a Z-score was computed from the unadjusted *p*-value using a two-tailed test conversion [Z = qnorm( *p*/2, lower.tail=FALSE) × sign(logFC)]. This calculation incorporates both the significance and direction of change, similar to the mouse dataset. The msigdbr was used to retrieve the mouse or human hallmark gene set as the input pathway ^75^. The output was filtered using minSize and maxSize to include only pathways with more than 15 genes but less than 500 to avoid noise, and for terms with an adjusted *p*-value less than 0.1.

### Gene correlation analysis

To quantify pairwise transcriptional similarity between datasets, we computed the overlap coefficient and Spearman correlations on differentially expressed genes (DEGs) between pairs of datasets. Both DEG lists were converted and/or subset to human nomenclature by mapping using Ensembl gene IDs corresponding to GRCh38.p14 (human) and GRCm39 (mouse) and retaining only one-to-one ortholog pairs to ensure direct comparability. For the overlap coefficient, we took significantly up and downregulated DEGs (adjusted p-value is less than or equal to 0.05, and the absolute logFC is greater than or equal to a 1.5 fold change) separately and calculated the intersection between the datasets divided by the size of the smaller set. Spearman correlation was computed on the logFC values across the union of all genes that were differentially expressed in at least one dataset and measured in both datasets. Statistical significance of correlations was determined using two-tailed tests, with p < 0.05 considered significant.

### Gene scores

The following gene lists were used to calculate gene scores using scanpy functions ^9,31,76^. Mouse/human nomenclature was interchanged by mapping the gene orthologs using GRCh38.p14 and GRCm39.

Cell cycle scoring: We downloaded the Regev lab cell cycle genes, as used in Seurat and the scverse tutorials, and converted the genes to mouse nomenclature using biomart (https://github.com/scverse/scanpy_usage/tree/master/180209_cell_cycle).

<u>Pandey:</u> *GADD45B, CAMK2D, CNTN1, VIM, NUPR1, SLC3A2, FOS, JUNB, JUN, DDIT3, KLK6, CD63, HMGB2, DYNLL1, CSF1, B2M, FBL, KLF6*

<u>IRAS1:</u> *CCL2, CCL20, CSF3, CXCL1, CXCL2, CXCL3, CXCL5, FTH1, PDCD5, PDLIM4, PITX3, PSME2, SERPINE1, SOD2, TNFAIP6*

<u>IRAS2:</u> *B2M, CALR, CARHSP1, CCL2, CD47, FTH1, ICAM1, LAP3, NFKBIA, NNMT, PDCD5, PFN1, PKM, PRELID1, PSME2, SOD2, TMSB10, VCAM1*

<u>Common innate immunity:</u> *IFI47, IIGP1, MDFIC, TOP2A, DYNLT1A, LGALS1, CXCL10, OASL1, ZBP1, DHX58, TRIM25, RSAD2, SLFN9, OAS1A, IFITM2, BCL3, IFI27L2A, OASL2, IRF7, IFIT1, BST2, ISG15, IFIT3B, IFIT3, IFITM3, RTP4*

<u>Remyelination DAO signature</u>: *COL6A1, GADD45G, COL27A1, COL6A2, NAV3, SHISA8, TMEM176A, CD63, COL13A1, FGFRL1, SNX10, PARP3, CXCL14, NRTN, SOCS3, ANXA2, TNFRSF1A, BCL3, B2M, PLVAP, GADD45B, SEMA4F*

<u>CPZ DAO signature:</u> *SPRY2, SYT4, CHCHD10, DTHD1, TRIB3, EPHA5, NUPR1, ATF4, CYB5R2, DDIT3, GDF15, CD44, CEBPG, VEGFA, CDH6, SLC7A5, S100A6, CDKN1A, RGS20, LMNA, VIM, EMP1, CLCF1, VGF, SERPINE1, RGS17, LGALS3, EGR1, IL6R*

<u>Hypoxia signature:</u> *B3GALT6, FOSL2, GLRX, NDRG1, SDC2, SDC3, PAM, PGM1, PLIN2, VEGFA, ENO2, ISG20, NFIL3, PIM1, PLAC8, TGFBI, RRAGD, BHLHE40, CHST2, CXCR4, LOX, PLAUR, SLC2A1, MAFF, CDKN1A, HEXA, MT1F*

<u>Inflammatory response signature:</u> *ABCA1, IL18R1, SLC31A2, IL10, IL15, IL1R1, IL4R, MXD1, NAMPT, INHBA, LDLR, NOD2, SLC31A1, OSM, OLR1, IFNAR1, SLC31A2, PTGER4, PLAUR, IL1B, MMP14, PTAFR, CHST2, AXL, CXCL9, CD14, CD69, CCL5, LDLR, P2RX4, CDKN1A, CD48*

### Meta-Neighborhood analysis

To determine cluster similarity in our cross-species analysis, we used MetaNeighbor 77. We merged the human-mouse datasets, and calculated the top 2000 HVGs for the oligodendrocyte or microglia lineage separately. The data was then passed to MetaNeighborUS using the fast version, with organism as study_id, and the broad cell type annotation as the ct_col variable.

## Notes

### Competing Interest Statement

The authors have declared no competing interest.

### Summary of Updates

All figures revised, supplemental files updated, new supplemental figures, author list updated, and rewritten text.

