## Supplementary Figures for "A comparative transcriptomic analysis of mouse demyelination models and Multiple Sclerosis lesions"

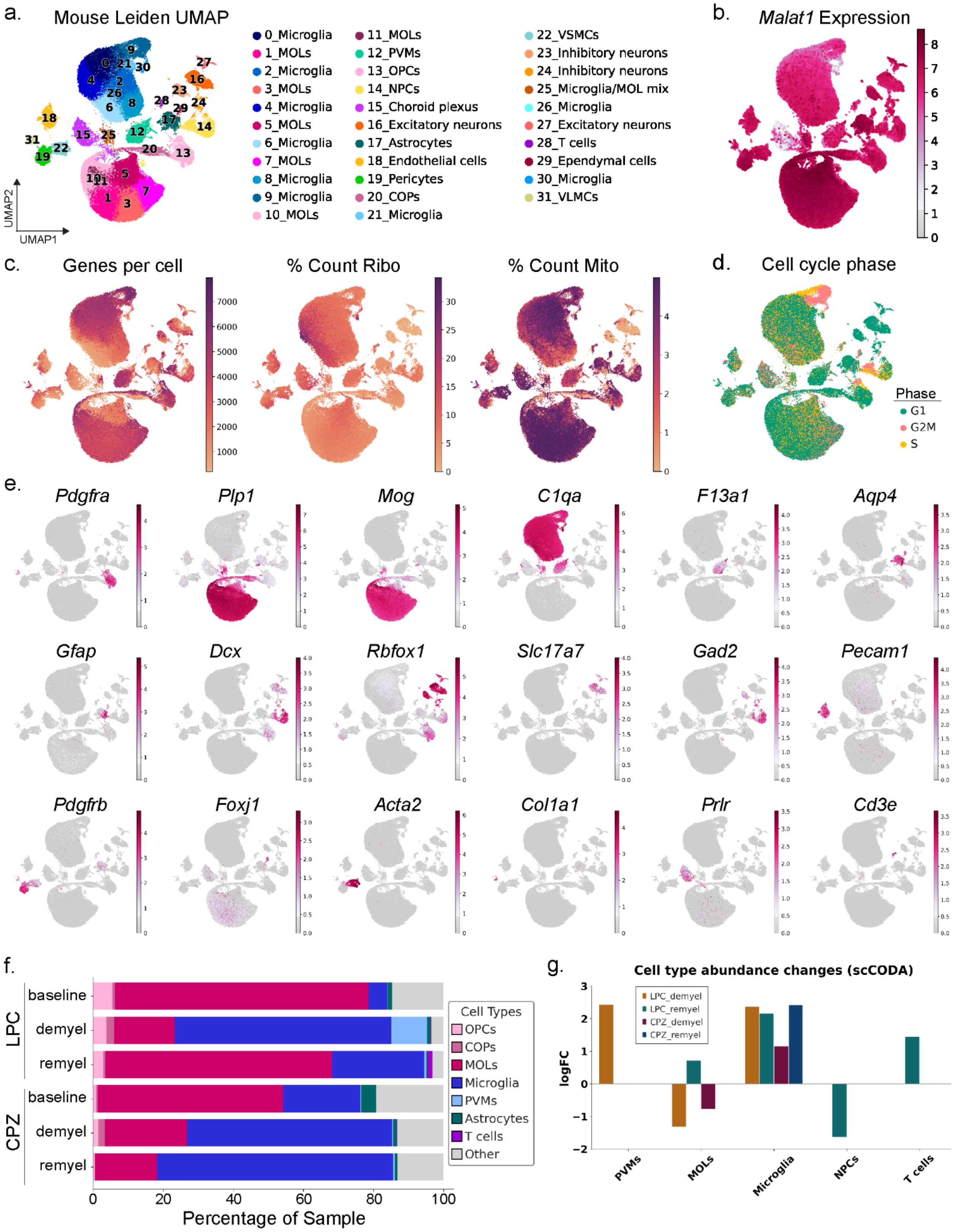

a.
Mouse Leiden UMAP
0_Microglia
1_MOLs
2_Microglia
3_MOLs
4_Microglia
5_MOLs
6_Microglia
7_MOLs
8_Microglia
9_Microglia
10_MOLs
11_MOLs
12_PVMs
13_OPCs
14_NPCs
15_Choroid plexus
16_Excitatory neurons
17_Astrocytes
18_Endothelial cells
19_Pericytes
20_COPs
21_Microglia
22_VSMCs
23_Inhibitory neurons
24_Inhibitory neurons
25_Microglia/MOL mix
26_Microglia
27_Excitatory neurons
28_T cells
29_Ependymal cells
30_Microglia
31_VLMCs
UMAP1
UMAP2
b.
Malat1 Expression
c.
Genes per cell
% Count Ribo
% Count Mito
d.
Cell cycle phase
Phase
G1
G2M
S
e.
Pdgfra
Plp1
Mog
C1qa
F13a1
Aqp4
Gfap
Dcx
Rbfox1
Slc17a7
Gad2
Pecam1
Pdgfrb
Foxj1
Acta2
Col1a1
Prlr
Cd3e
f.
LPC
baseline
demyel
remyel
CPZ
baseline
demyel
remyel
Cell Types
OPCs
COPs
MOLs
Microglia
PVMs
Astrocytes
T cells
Other
Percentage of Sample
g.
Cell type abundance changes (scCODA)
LPC_demyel
LPC_remyel
CPZ_demyel
CPZ_remyel
logFC
PVMs
MOLs
Microglia
NPCs
T cells

**Supplementary Figure 1: Quality control of mouse data.**

a. UMAP of the fully integrated mouse dataset, with cell types labeled on the leiden clustering.
b. UMAP showing *Malat1* expression, where low expression serves as a proxy for low-quality cells.
c. UMAPs displaying key quality control metrics, including genes per cell, percentage of ribosomal counts (% Count Ribo), and percentage of mitochondrial counts (% Count Mito).
d. Cell cycle phase analysis plotted on the integrated UMAP. Three actively dividing cell populations are identified, corresponding to OPCs, microglia, and NPCs.
e. UMAPs showing expression of the indicated marker genes, corresponding to the dot plot in Figure 1c.
f. Stacked bar chart showing the distribution of annotated cell types across each timepoint. Corresponding metadata is available in Supplemental Data 1.
g. Bar chart depicting the significant cell compositional changes calculated by scCODA. Calculations were made per condition and compared to the respective baseline. Pericytes were used as the reference cell type.

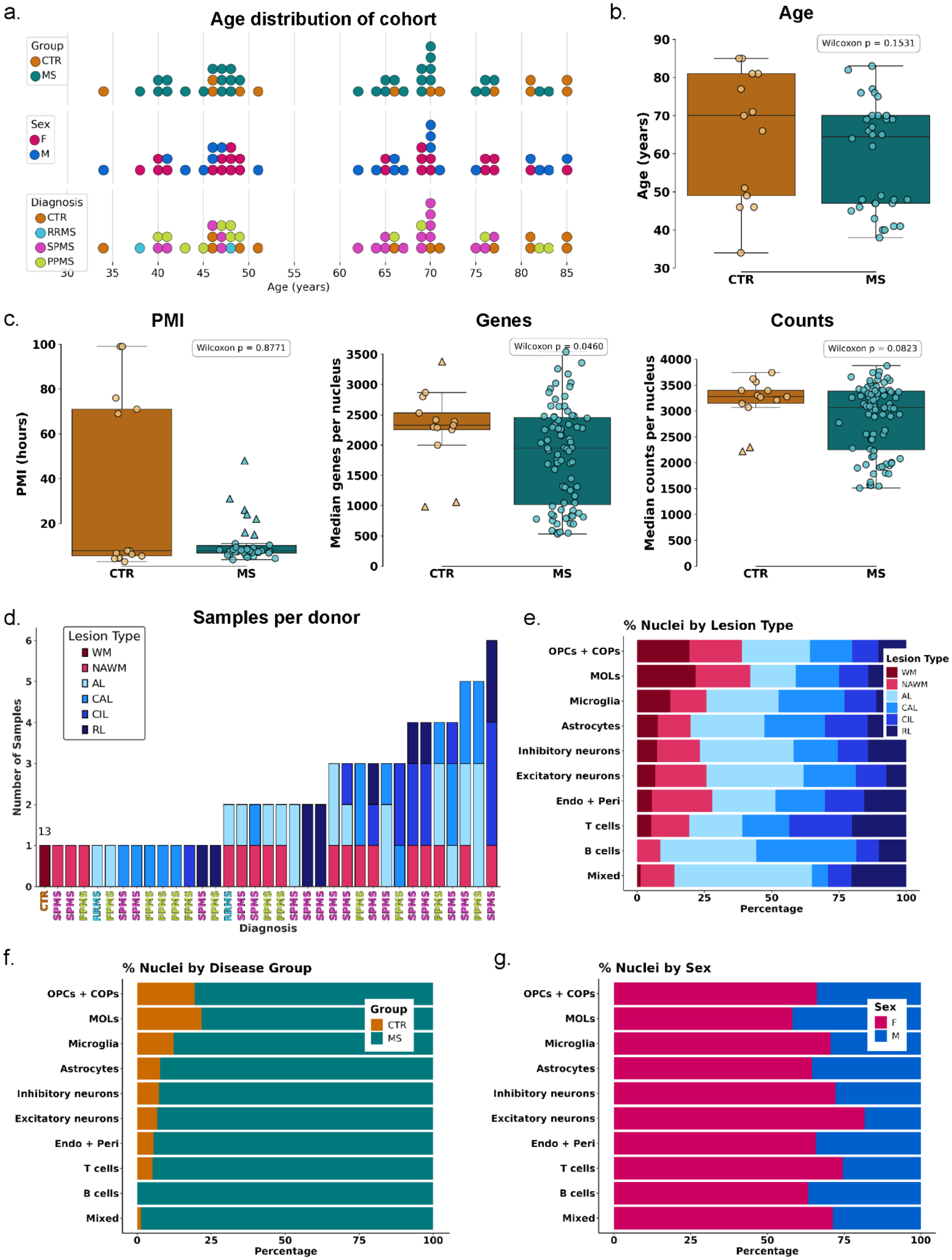

a.
Age distribution of cohort
Group
CTR
MS
Sex
F
M
Diagnosis
CTR
RRMS
SPMS
PPMS
Age (years)
b.
Age
Wilcoxon p = 0.1531
Age (years)
CTR
MS
c.
PMI
Wilcoxon p = 0.8771
PMI (hours)
Genes
Wilcoxon p = 0.0460
Median genes per nucleus
Counts
Wilcoxon p = 0.0823
Median counts per nucleus
d.
Samples per donor
Lesion Type
WM
NAWM
AL
CAL
CIL
RL
Number of Samples
13
Diagnosis
e.
% Nuclei by Lesion Type
OPCs + COPs
MOLs
Microglia
Astrocytes
Inhibitory neurons
Excitatory neurons
Endo + Peri
T cells
B cells
Mixed
Percentage
f.
% Nuclei by Disease Group
Group
CTR
MS
Percentage
g.
% Nuclei by Sex
Sex
F
M
Percentage

**Supplementary Figure 2: Statistical comparison and quality control of human metadata.**

a. Age distribution of patient cohort, colored by group (control (CTR) and MS patients, top), sex (female (F), male (M), middle) and diagnosis (relapse-remitting MS (RRMS), secondary progressive MS (SPMS) and primary progressive MS (PPMS), bottom).
b. Box plot showing the age distribution of CTR and MS patient donors. As the data are not normally distributed (confirmed using the Shapiro-Wilk test), a Mann-Whitney U test was used to compare the groups and indicated no significant difference. The boxes represent the interquartile range (IQR), with the center line showing the median. Whiskers extend to the most extreme data points within 1.5 × IQR from the lower (Q1) and upper (Q3) quartiles. All individual donor ages are shown as points.
c. Box plot (as in b) showing the post-mortem interval (PMI), median number of genes per nucleus, and median counts per nucleus across donor samples in CTR and MS groups. Outliers, identified using the IQR method, are shown as triangles. As the data was not normally distributed, a Mann-Whitney U test was used. A statistically significant difference was observed between groups for PMI and genes per nucleus.
d. Stacked bar chart showing the number of unique samples per donor. Each bar represents one patient and is colored by lesion type, except for the 13 controls (Ctr) represented by 1 bar. The x-axis shows the patient diagnoses (SP: secondary progressive, PP: primary progressive, RR: relapse remitting). For MS patients, thirteen individuals contributed one sample and 21 individuals contributed more than one sample.
e. Stacked barplot showing the percentage contribution to the respective cell type cluster color by lesion type.
f. Stacked barplot showing the percentage contribution to the respective cell type cluster colored by group.
g. Stacked barplot showing the percentage contribution to the respective cell type cluster colored by sex.

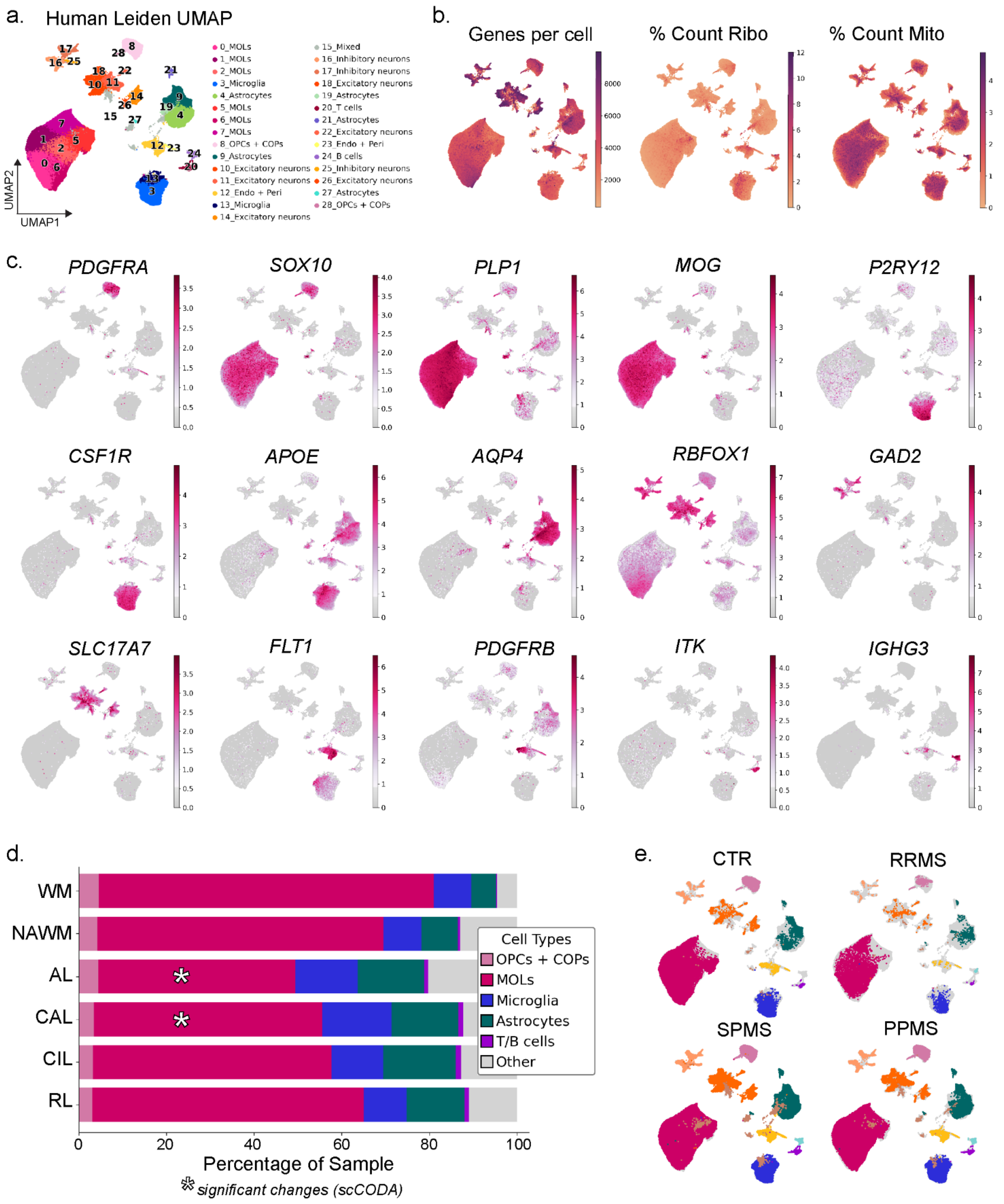


**Supplementary Figure 3: Quality control of integration from human samples.**

a. UMAP of the integrated dataset with cell types annotated on the leiden clustering.

b. UMAPs displaying key quality control metrics, including genes per nucleus, percentage of ribosomal counts (% Count Ribo), and percentage of mitochondrial counts (% Count Mito).
c. UMAPs showing expression of the indicated marker genes, corresponding to the dot plot in Figure 1g.
d. Stacked bar chart showing the proportion of glial cell types per lesion type. (Other = neurons, Endo + Peri, and mixed cell types). Asterisks denote significant compositional changes calculated using scCODA by comparing each lesion type to WM and using the cluster “Endo + Peri” as the reference cell type
e. UMAPs separated by disease status, (Ctr: control, RRMS: relapse remitting MS, SPMS: secondary progressive MS, PPMS: primary progressive MS).

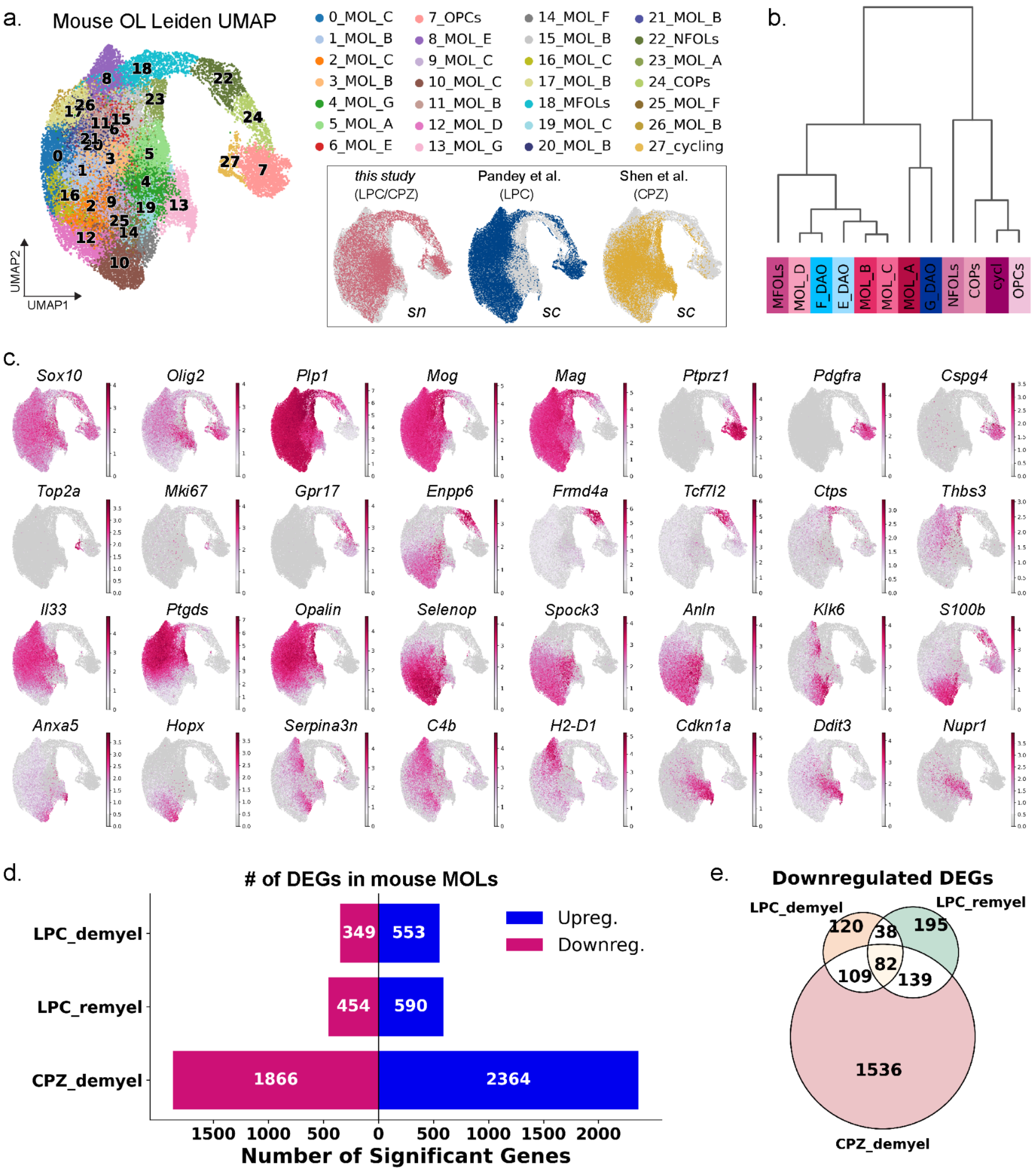


**Supplementary Figure 4: Mouse oligodendrocyte lineage clustering, marker expression, and differential expression analysis.**

a. UMAP of the OL subset with cell types labeled based on leiden clustering. Inset UMAPs show samples separated and colored by dataset origin, labeled by the demyelination treatment and sequencing type.

b. Dendrogram showing the hierarchical clustering of the cell types, colored as the cell types in the UMAP in Figure 2a.
c. UMAPs showing expression of the indicated marker genes, corresponding to the dot plot in Figure 2b.
d. Stacked bar chart showing the upregulated (blue) and downregulated (magenta) differentially expressed genes (DEGs) in the indicated cell type and timepoint.
e. Venn diagram illustrating the unique and overlapping significantly downregulated genes (logFC ≤ 1.5, adjusted $p < 0.05$) calculated from the pseudobulk of all MOLs in LPC or CPZ at demyelination, and LPC at remyelination.

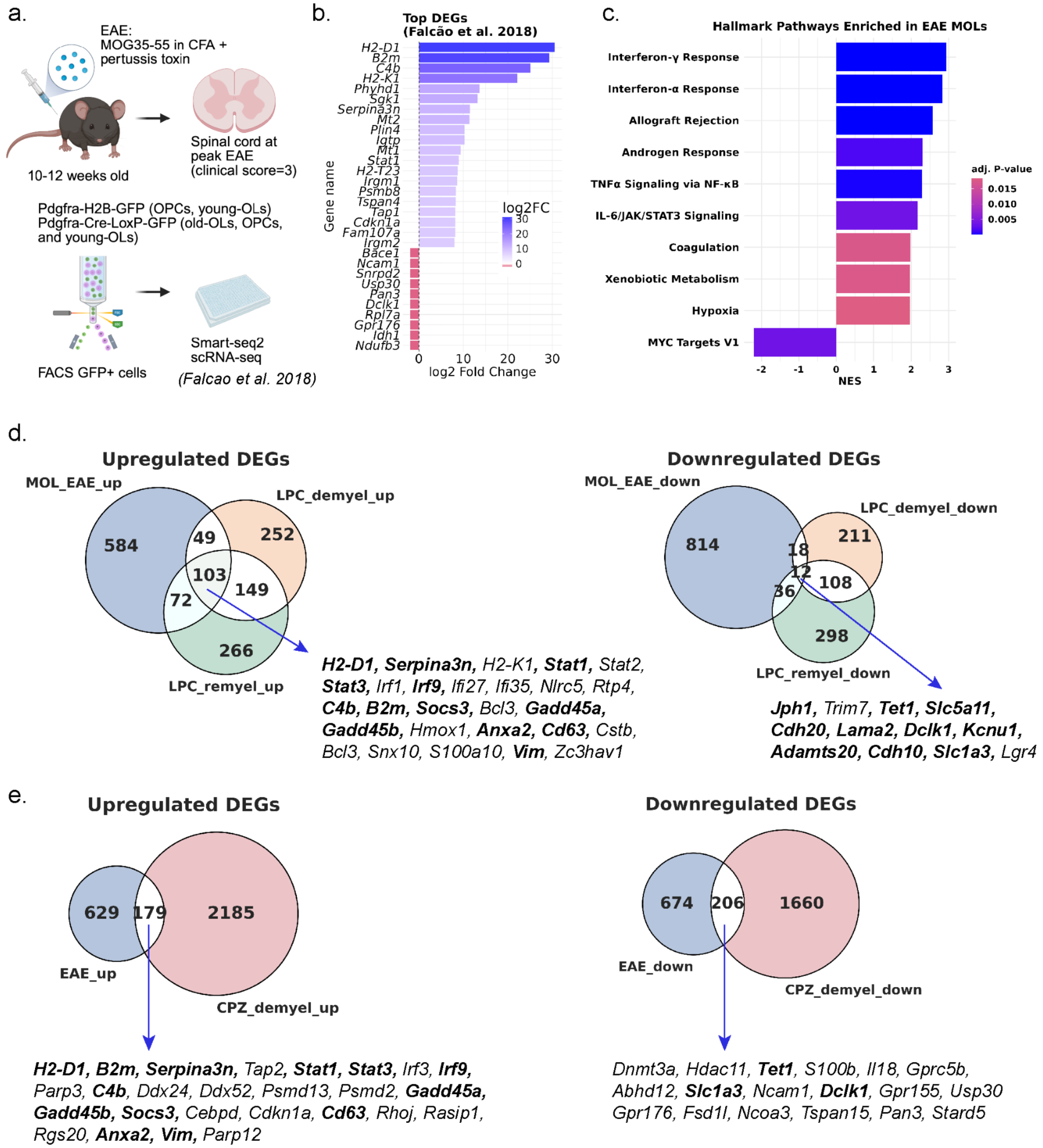


**Supplementary Figure 5: Comparison of LPC and CPZ gene expression changes with a mouse EAE model.**

a. Schematic of the EAE dataset used for comparison. EAE was induced in transgenic adult mice that label OPCs and OLs with GFP. At the peak of the disease, mice were sacrificed and spinal cord OPCs and OLs were FACS sorted. Single cell RNA-seq data was collected using a plate-based protocol called Smart-seq2.

b. Bar chart of the top up and downregulated genes from the EAE dataset. The provided supplementary table of DEGs, calculated using MAST, includes only significant DEGs with a logFC of at least 1.5 fold.
c. Bar plot showing the enrichment of Hallmark pathways in the EAE dataset. Ranking score was set to the logFC, as this is the only available data for the DEG list provided from the original manuscript. Color indicates the adjusted *p*-value and bar length is the normalized enrichment score (NES).
d. Overlap of upregulated (left) and downregulated (right) genes between EAE and LPC treated oligodendrocytes. A subset of genes is listed from the intersection of all 3 datasets. Bolded genes are shared between LPC, CPZ and EAE.
e. Overlap of upregulated (left) and downregulated (right) genes between EAE and CPZ treated oligodendrocytes.A subset of genes is listed from the intersection of all 3 datasets. Bolded genes are shared between LPC, CPZ and EAE.

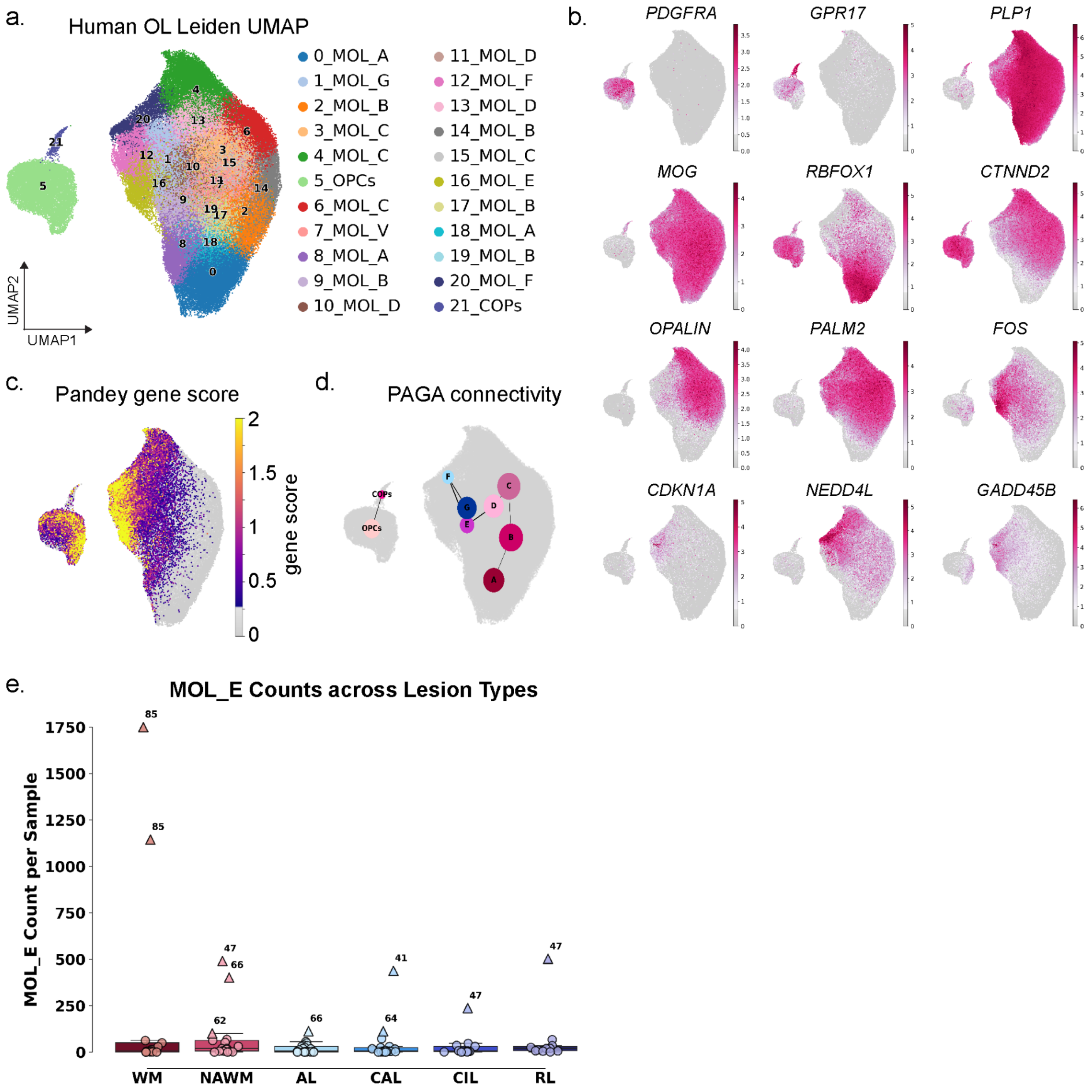


**Supplementary Figure 6: Human oligodendrocyte lineage clustering, marker expression and QC.**

a. UMAP of the human oligodendrocyte lineage with cell types labeled on the leiden clustering.
b. UMAPs showing expression of the indicated marker genes, corresponding to the dot plot in Figure 4b.
c. Gene score calculated using the expression of DAO genes identified by Pandey et. al. in mouse models that are conserved in MS patient samples (genes listed in Methods).
d. UMAP showing PAGA connectivity with edge weights above 0.25 between the annotated cell types as shown in Figure 4a.

e. Box plots of MOL_E cell type count across lesion types, with each dot representing a single sample. Triangles indicate samples identified as statistical outliers based on interquartile range (IQR), defined as samples exceeding 1.5× IQR above the third quartile. Outliers are annotated with the age at death of the donor in years.

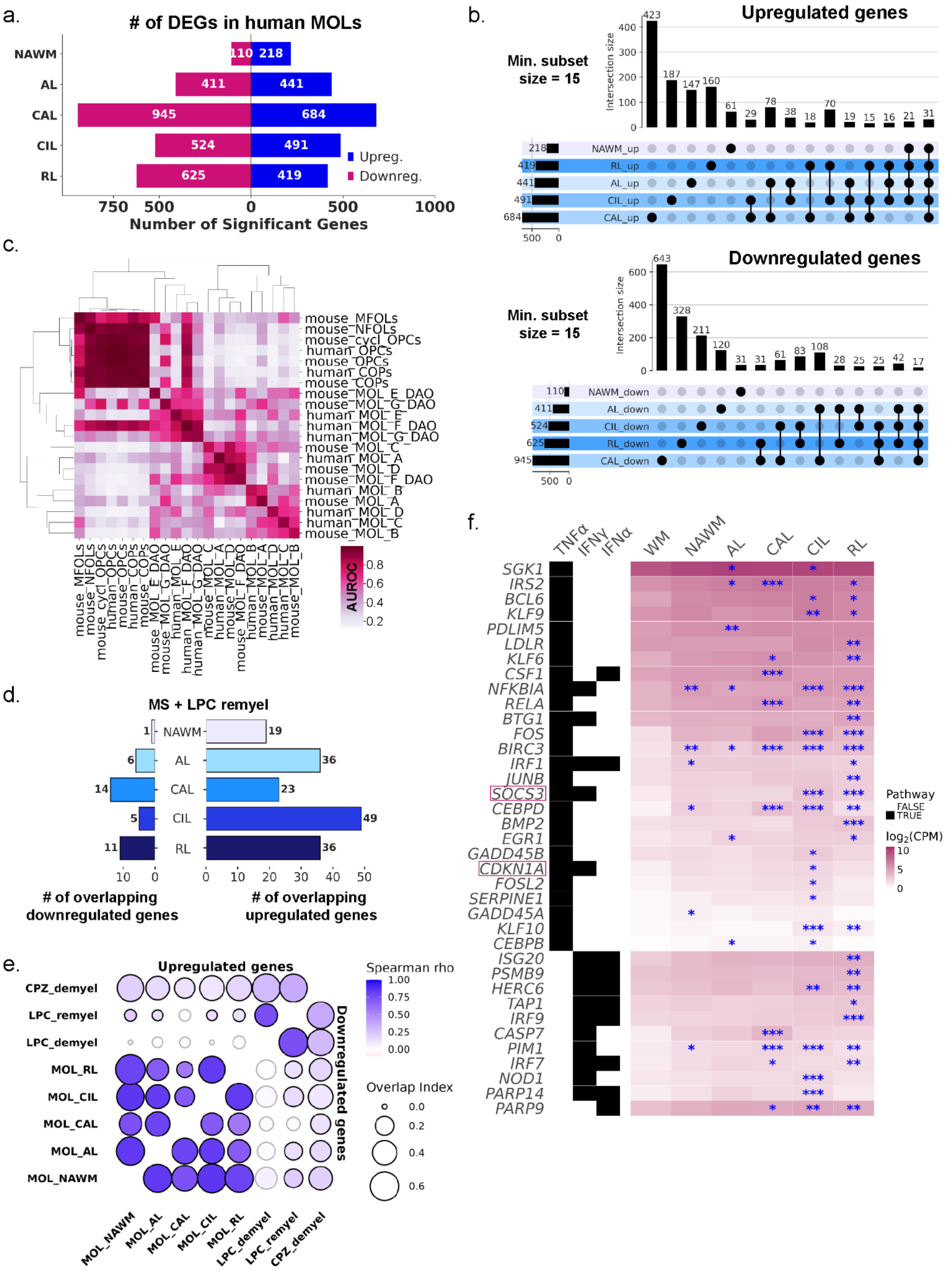
a.
### of DEGs in human MOLs
NAWM
AL
CAL
CIL
RL
110
218
411
441
945
684
524
491
625
419
Upreg.
Downreg.
750
500
0
500
1000
Number of Significant Genes
b.
Upregulated genes
Min. subset size = 15
Intersection size
423
187
147
160
61
29
78
38
18
70
19
15
16
21
31
NAWM_up
RL_up
AL_up
CIL_up
CAL_up
218
419
441
491
684
Downregulated genes
Min. subset size = 15
643
328
211
120
31
31
61
83
108
28
25
25
42
17
NAWM_down
AL_down
CIL_down
RL_down
CAL_down
110
411
524
625
945
c.
mouse_MFOLs
mouse_NFOLs
mouse_cycl_OPCs
human_OPCs
mouse_OPCs
human_COPs
mouse_COPs
mouse_MOL_E_DAO
mouse_MOL_G_DAO
human_MOL_E
human_MOL_F_DAO
human_MOL_G_DAO
mouse_MOL_C
human_MOL_A
mouse_MOL_D
mouse_MOL_F_DAO
human_MOL_B
mouse_MOL_A
human_MOL_D
human_MOL_C
mouse_MOL_B
AUROC
0.8
0.6
0.4
0.2
d.
MS + LPC remyel
NAWM
AL
CAL
CIL
RL
1
6
14
5
11
19
36
23
49
36
### of overlapping downregulated genes
### of overlapping upregulated genes
e.
Upregulated genes
Downregulated genes
CPZ_demyel
LPC_remyel
LPC_demyel
MOL_RL
MOL_CIL
MOL_CAL
MOL_AL
MOL_NAWM
Spearman rho
1.00
0.75
0.50
0.25
0.00
Overlap Index
0.0
0.2
0.4
0.6
MOL_NAWM
MOL_AL
MOL_CAL
MOL_CIL
MOL_RL
LPC_demyel
LPC_remyel
CPZ_demyel
f.
TNFα
IFNγ
IFNα
WM
NAWM
AL
CAL
CIL
RL
SGK1
IRS2
BCL6
KLF9
PDLIM5
LDLR
KLF6
CSF1
NFKBIA
RELA
BTG1
FOS
BIRC3
IRF1
JUNB
SOCS3
CEBPD
BMP2
EGR1
GADD45B
CDKN1A
FOSL2
SERPINE1
GADD45A
KLF10
CEBPB
ISG20
PSMB9
HERC6
TAP1
IRF9
CASP7
PIM1
IRF7
NOD1
PARP14
PARP9
Pathway
FALSE
TRUE
log2(CPM)
10
5
0

**Supplementary Figure 7: Human oligodendrocyte DEG analysis and comparison with mouse models.**

a. Stacked bar chart showing the total number of upregulated (blue) and downregulated (magenta) DEGs across all MOLs per lesion type.
b. Upset plots showing the unique (single points) and overlapping (connected points) DEGs in MOLs across lesion conditions that are upregulated (logFC ≥ 1.5, padj < 0.05, top) and downregulated ((logFC ≤ 1.5, padj < 0.05, bottom). Overlap size was filtered to include only points with 15 or more shared genes.
c. Heatmap of AUROC scores between mouse and human OL lineage cells calculated using MetaNeighbor analysis. Dendrograms were generated by hierarchical clustering of Euclidean distances using average linkage.
d. Bar chart showing the overlap of upregulated and downregulated DEGs shared between MOLs in the mouse LPC remyelination timepoint and the MS lesions.
e. Scatterplot matrix showing the pairwise similarity between datasets, with upper-left triangles showing concordance for upregulated DEGs and lower-right triangles for downregulated DEGs. Circle size indicates the fraction of shared DEGs (overlap coefficient). Fill color represents the Spearman correlation coefficient (ρ) of log fold-changes among shared genes and black outlines denote statistically significant correlations (p < 0.05).
f. Heatmap showing expression of genes in the indicated pathways across lesion conditions. Heatmaps are colored by the average $\log_2$-transformed CPM expression values. Gene–pathway associations were assigned to pathways using the Hallmark sets and are shown in the left annotation panel, where black tiles indicate pathway membership. Genes are hierarchically ordered first by pathway inclusion and then by average expression across all conditions. Blue asterisks indicate genes with significant upregulation (* for $p < 0.05$, ** for $p < 0.01$, *** for $p < 0.001$). (TNFα: tumor necrosis factor alpha, INFγ: interferon gamma, IFNα: interferon alpha).

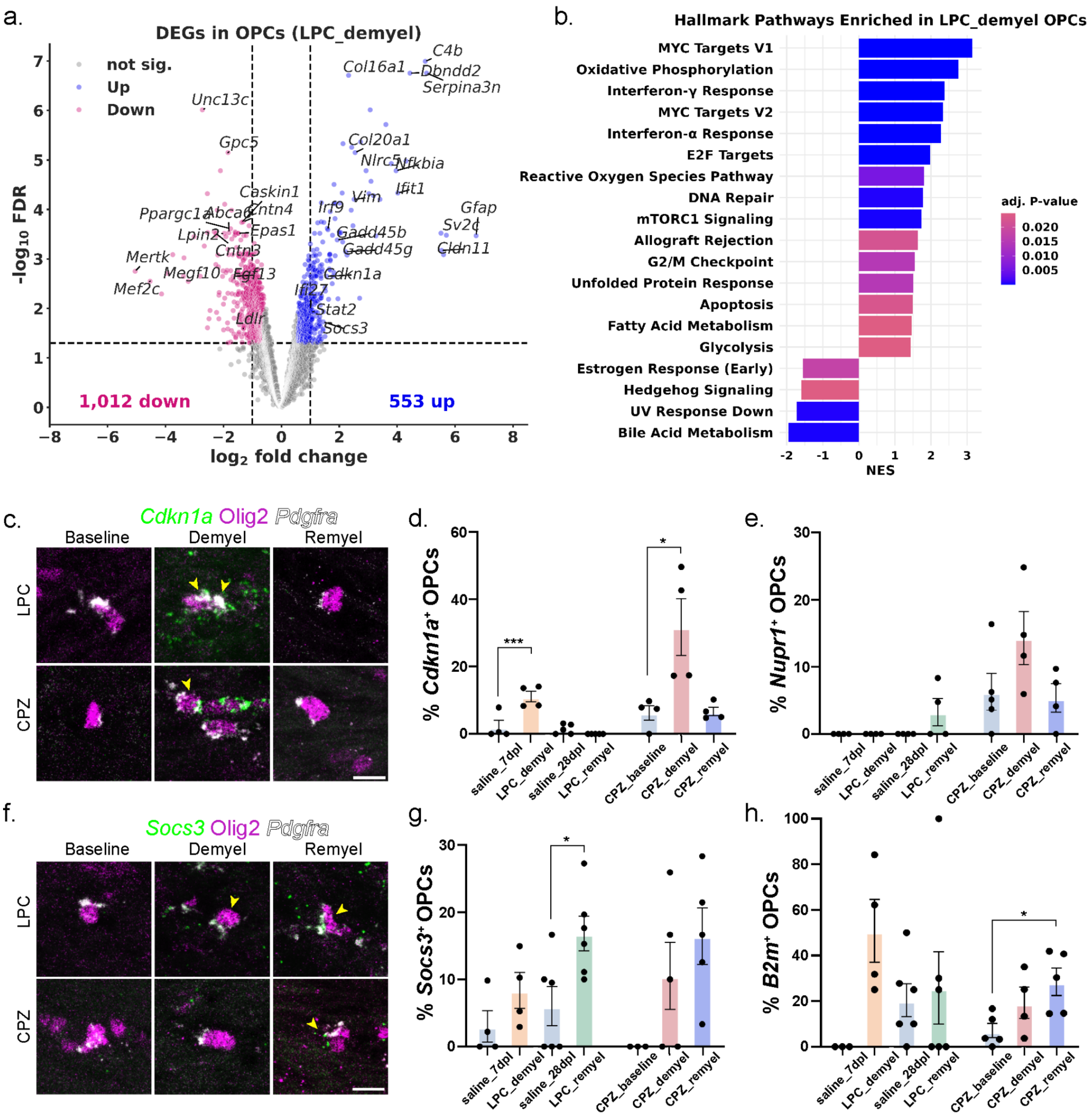


**Supplementary Figure 8: Differential gene expression in OPCs from LPC and CPZ mouse models.**

a. Volcano plot showing the statistical significance and logFC of all genes (each dot) tested for differential expression in OPCs following demyelination by LPC treatment. Significantly upregulated genes are colored blue, and downregulated genes are colored magenta. Dashed horizontal and vertical lines indicate the significance thresholds for *padj* and logFC, respectively.
b. Bar plot showing the enrichment of Hallmark pathways from OPC gene expression at LPC demyelination. Color indicates the adjusted *p*-value and bar length is the normalized enrichment score (NES).

c. Representative RNAscope images from the mouse corpus callosum showing *Cdkn1a* expression. Yellow arrows show *Cdkn1a*$^{+}$ OPCs. Time points for LPC are: baseline=saline injected at 7dpl, demyel= LPC injected at 7dpl, and remyel= LPC injected at 28dpl; and for CPZ: baseline= normal diet, demyel= 5 weeks of CPZ chow, and remyel= 5 weeks CPZ chow and 3 weeks of recovery. (Scale bar = 10 μm).
d. Bar chart showing the percentage of *Cdkn1a*$^{+}$ OPCs (*Cdkn1a*$^{+}$Olig2$^{+}$*Pdgfra*$^{+}$ / Olig2$^{+}$*Pdgfra*$^{+}$) across lesion conditions. Bars represent group means, dots show individual animals, and error bars indicate SEM. Statistical significance was assessed by one-way ANOVA with Tukey's multiple comparisons test (***$p$ = 0.0006, *$p$ = 0.0154*).*
e. Bar chart showing the percentage of *Nupr1*$^{+}$ OPCs (*Nupr1*$^{+}$Olig2$^{+}$*Pdgfra*$^{+}$ / Olig2$^{+}$*Pdgfra*$^{+}$) across lesion conditions. Bars represent group means, dots show individual animals, and error bars indicate SEM.
f. Representative RNAscope images from the mouse corpus callosum showing *Socs3* expression. Yellow arrows show *Socs3*$^{+}$ OPCs. Time points are the same as above.
g. Bar chart showing the percentage of *Socs3*$^{+}$ OPCs (*Socs3*$^{+}$Olig2$^{+}$*Pdgfra*$^{+}$ / Olig2$^{+}$*Pdgfra*$^{+}$) across lesion conditions. Bars represent group means, dots show individual animals, and error bars indicate SEM. Statistical significance was assessed by one-way ANOVA with Tukey's multiple comparisons test (*$p$ = 0.0347*).*
h. Bar chart showing the percentage of *B2m*$^{+}$ OPCs (*B2m*$^{+}$Olig2$^{+}$*Pdgfra*$^{+}$ / Olig2$^{+}$*Pdgfra*$^{+}$) across lesion conditions. Bars represent group means, dots show individual animals, and error bars indicate SEM. Statistical significance was assessed by one-way ANOVA with Tukey's multiple comparisons test (*$p$ = 0.0347*).*

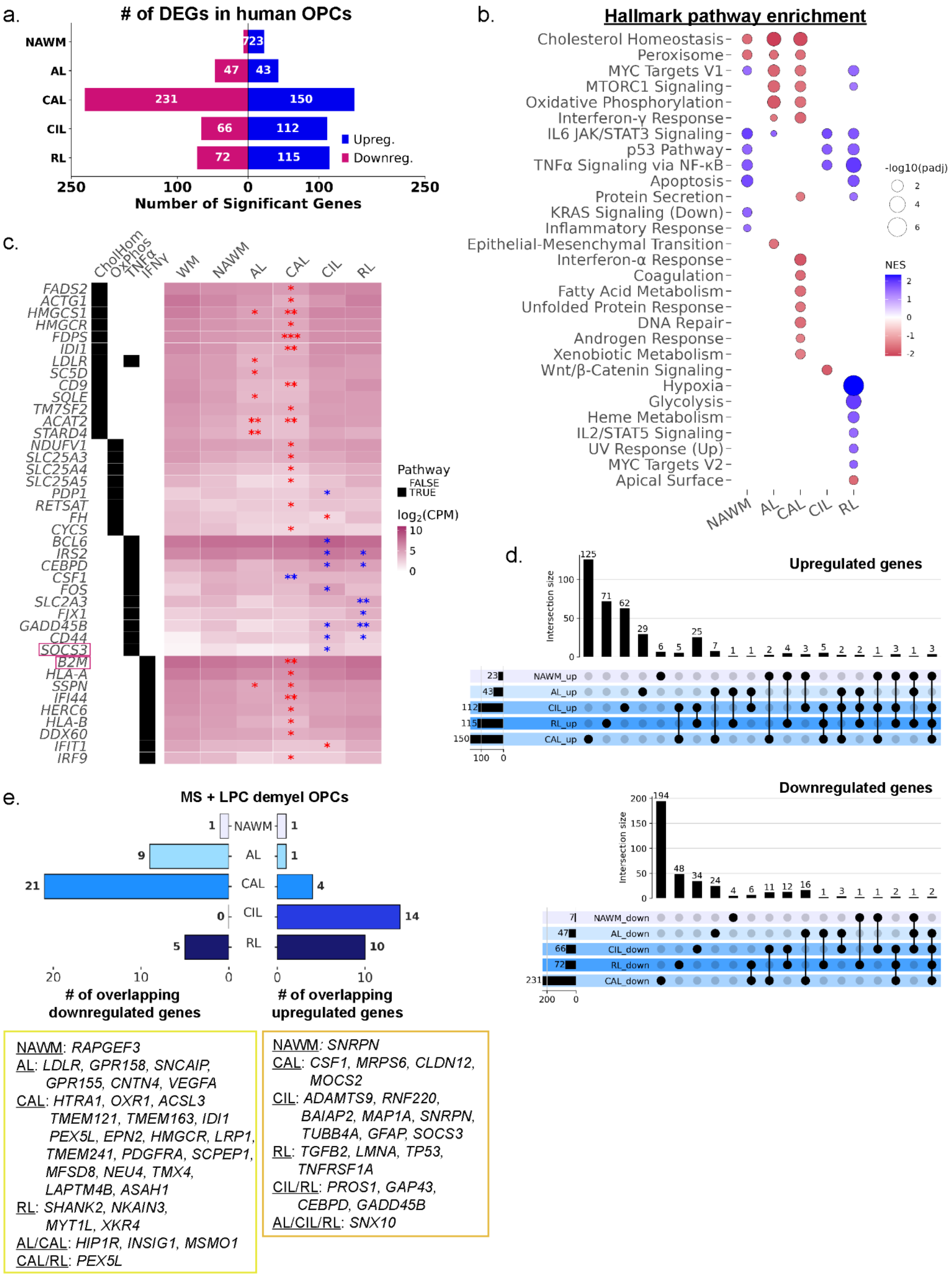

a.
### of DEGs in human OPCs
NAWM
AL
CAL
CIL
RL
7
23
47
43
231
150
66
112
72
115
Upreg.
Downreg.
250
100
0
100
250
Number of Significant Genes
b.
Hallmark pathway enrichment
Cholesterol Homeostasis
Peroxisome
MYC Targets V1
MTORC1 Signaling
Oxidative Phosphorylation
Interferon-γ Response
IL6 JAK/STAT3 Signaling
p53 Pathway
TNFα Signaling via NF-κB
Apoptosis
Protein Secretion
KRAS Signaling (Down)
Inflammatory Response
Epithelial-Mesenchymal Transition
Interferon-α Response
Coagulation
Fatty Acid Metabolism
Unfolded Protein Response
DNA Repair
Androgen Response
Xenobiotic Metabolism
Wnt/β-Catenin Signaling
Hypoxia
Glycolysis
Heme Metabolism
IL2/STAT5 Signaling
UV Response (Up)
MYC Targets V2
Apical Surface
NAWM
AL
CAL
CIL
RL
-log10(padj)
2
4
6
NES
2
1
0
-1
-2
c.
CholHom
OxPhos
TNFα
IFNγ
WM
NAWM
AL
CAL
CIL
RL
FADS2
ACTG1
HMGCS1
HMGCR
FDPS
IDI1
LDLR
SC5D
CD9
SQLE
TM7SF2
ACAT2
STARD4
NDUFV1
SLC25A3
SLC25A4
SLC25A5
PDP1
RETSAT
FH
CYCS
BCL6
IRS2
CEBPD
CSF1
FOS
SLC2A3
FJX1
GADD45B
CD44
SOCS3
B2M
HLA-A
SSPN
IFI44
HERC6
HLA-B
DDX60
IFIT1
IRF9
Pathway
FALSE
TRUE
log2(CPM)
10
5
0
d.
Upregulated genes
Intersection size
125
71
62
29
6
5
25
7
1
1
2
4
3
5
2
2
1
3
1
3
NAWM_up
AL_up
CIL_up
RL_up
CAL_up
23
43
112
115
150
100
0
Downregulated genes
194
48
34
24
4
6
11
12
16
1
3
1
1
2
1
2
NAWM_down
AL_down
CIL_down
RL_down
CAL_down
7
47
66
72
231
200
0
e.
MS + LPC demyel OPCs
NAWM
AL
CAL
CIL
RL
1
9
21
0
5
1
1
4
14
10
20
10
0
0
10
### of overlapping downregulated genes
### of overlapping upregulated genes
NAWM: RAPGEF3
AL: LDLR, GPR158, SNCAIP, GPR155, CNTN4, VEGFA
CAL: HTRA1, OXR1, ACSL3 TMEM121, TMEM163, IDI1 PEX5L, EPN2, HMGCR, LRP1, TMEM241, PDGFRA, SCPEP1, MFSD8, NEU4, TMX4, LAPTM4B, ASAH1
RL: SHANK2, NKAIN3, MYT1L, XKR4
AL/CAL: HIP1R, INSIG1, MSMO1
CAL/RL: PEX5L
NAWM: SNRPN
CAL: CSF1, MRPS6, CLDN12, MOCS2
CIL: ADAMTS9, RNF220, BAIAP2, MAP1A, SNRPN, TUBB4A, GFAP, SOCS3
RL: TGFB2, LMNA, TP53, TNFRSF1A
CIL/RL: PROS1, GAP43, CEBPD, GADD45B
AL/CIL/RL: SNX10

**Supplementary Figure 9: OPC differential gene expression in humans and comparison between species.**

a. Stacked bar chart showing the total number of upregulated (blue) and downregulated (magenta) DEGs across OPCs in each lesion type.
b. Dot plot showing the enrichment of Hallmark pathways per lesion type in OPCs. Dot color indicates normalized enrichment score (NES), and size reflects –log10(adjusted *p*).
c. Heatmap showing the average $\log_2$-transformed CPM expression values of genes associated with the indicated pathways with significantly altered expression in at least one MS lesion condition in OPCs. Gene–pathway associations were assigned to pathways using the Hallmark sets and are shown in the left annotation panel, where black tiles indicate pathway membership. Genes are hierarchically ordered first by pathway inclusion and then by average expression across all conditions. Blue asterisks indicate genes with significant upregulation, and red asterisks indicated significant downregulation (* for $p < 0.05$, ** for $p < 0.01$, *** for $p < 0.001$). Full DEG results are reported in Supplementary table 7. (CholHom: cholesterol homeostasis, OxPhos: oxidative phosphorylation, INFγ: interferon gamma, TNFα: tumor necrosis factor alpha).
d. Upset plots showing the unique (single points) and overlapping (connected points) DEGs in MOLs across lesion conditions that are upregulated (logFC ≥ 1.5, padj < 0.05, top) and downregulated ((logFC ≤ 1.5, padj < 0.05, bottom).
e. Bar chart showing the overlap of upregulated (right) and downregulated (left) DEGs in OPCs between LPC at demyelination and the MS lesion types. Genes in boxes below the plots represent genes found in the unique overlap, or shared between mouse OPCs and multiple lesion types, as indicated.

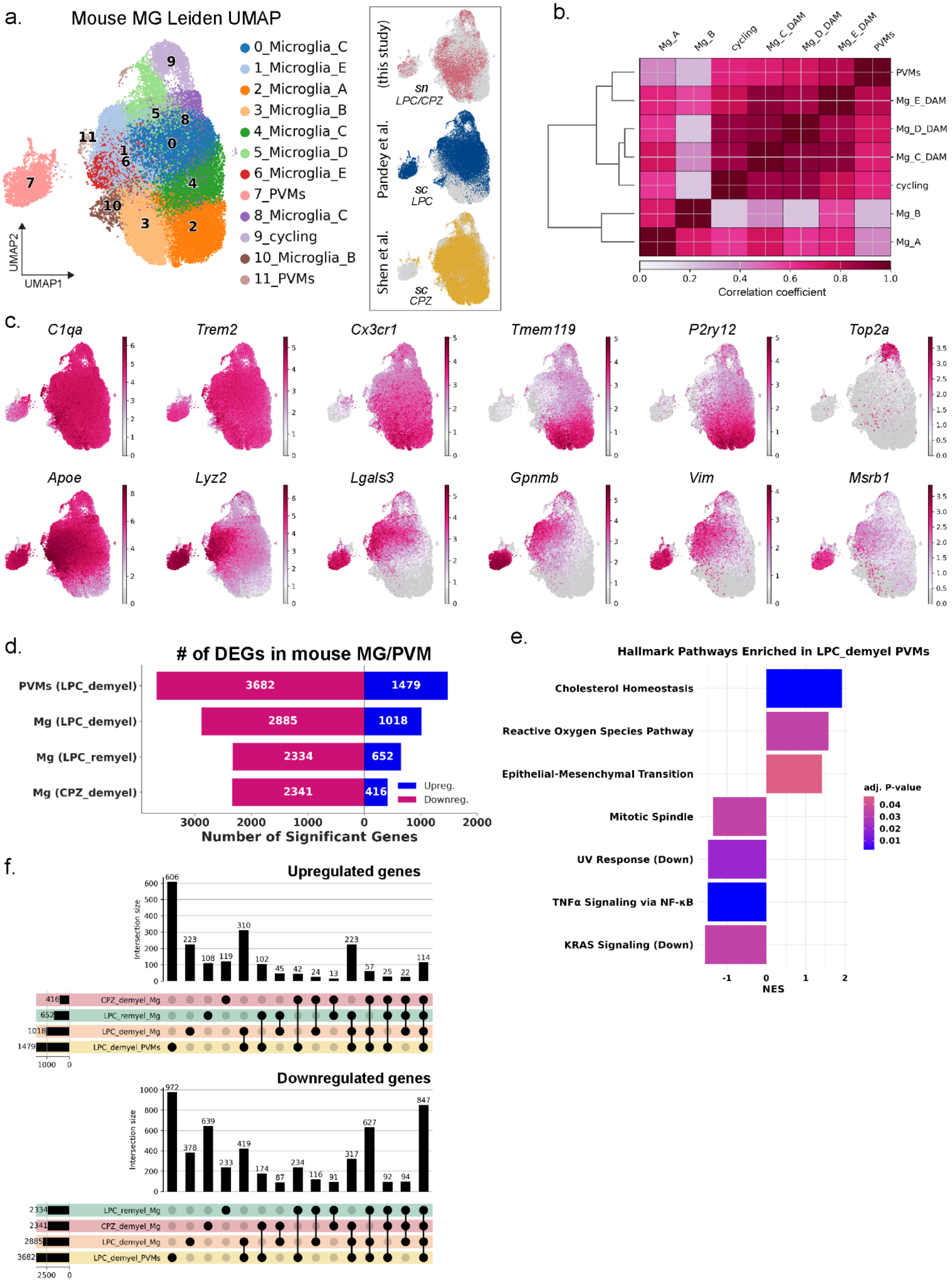
a.
Mouse MG Leiden UMAP
0_Microglia_C
1_Microglia_E
2_Microglia_A
3_Microglia_B
4_Microglia_C
5_Microglia_D
6_Microglia_E
7_PVMs
8_Microglia_C
9_cycling
10_Microglia_B
11_PVMs
UMAP1
UMAP2
(this study)
sn LPC/CPZ
Pandey et al.
sc LPC
Shen et al.
sc CPZ
b.
Mg_A
Mg_B
cycling
Mg_C_DAM
Mg_D_DAM
Mg_E_DAM
PVMs
Correlation coefficient
c.
C1qa
Trem2
Cx3cr1
Tmem119
P2ry12
Top2a
Apoe
Lyz2
Lgals3
Gpnmb
Vim
Msrb1
d.
### of DEGs in mouse MG/PVM
PVMs (LPC_demyel)
Mg (LPC_demyel)
Mg (LPC_remyel)
Mg (CPZ_demyel)
3682
1479
2885
1018
2334
652
2341
416
Upreg.
Downreg.
Number of Significant Genes
e.
Hallmark Pathways Enriched in LPC_demyel PVMs
Cholesterol Homeostasis
Reactive Oxygen Species Pathway
Epithelial-Mesenchymal Transition
Mitotic Spindle
UV Response (Down)
TNFα Signaling via NF-κB
KRAS Signaling (Down)
adj. P-value
NES
f.
Upregulated genes
Intersection size
CPZ_demyel_Mg
LPC_remyel_Mg
LPC_demyel_Mg
LPC_demyel_PVMs
Downregulated genes
Intersection size
LPC_remyel_Mg
CPZ_demyel_Mg
LPC_demyel_Mg
LPC_demyel_PVMs

**Supplementary Figure 10: Mouse microglia/PVM clustering, marker expression and differential expression analysis.**

a. UMAP of the MG and PVM subset with cell types labeled based on leiden clustering. Inset UMAPs show samples separated and colored by dataset origin, labeled by the demyelination treatment and sequencing type.
b. Correlation matrix and dendrogram showing the hierarchical clustering of the cell types.
c. UMAPs showing expression of the indicated marker genes, corresponding to the dot plot in Figure 5b.
d. Stacked bar chart showing the total number of upregulated (blue) and downregulated (magenta) DEGs in MG and PVMs compared to baseline per timepoint as indicated.
e. Bar plot showing the enrichment of Hallmark pathways from the gene expression changes in PVMs at LPC demyelination vs all microglia/PVM cells at LPC baseline. Color indicates the adjusted *p*-value and bar length is the normalized enrichment score (NES).
f. Upset plots showing the unique (single points) and overlapping (connected points) DEGs from PVM and microglia across lesion conditions that are upregulated (logFC $\geq$ 1.5, padj $< 0.05$, top) and downregulated ((logFC $\leq$ 1.5, padj $< 0.05$, bottom).

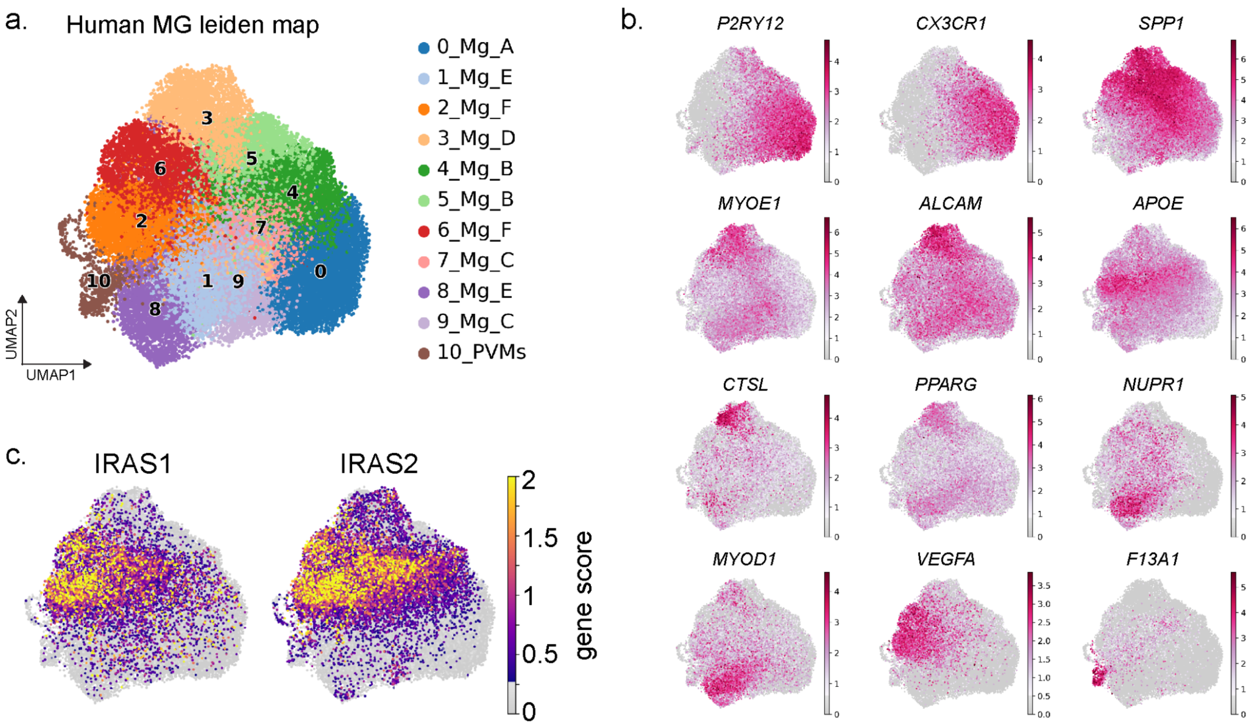


**Supplementary Figure 11: Human microglia lineage clustering, marker expression and QC.**

a. UMAP of the human MG and PVM subset with cell types labeled based on leiden clustering.
b. UMAPs showing expression of the indicated marker genes, corresponding to the dot plot in Figure 6b.
c. UMAPs of gene scores based on expression of inflammatory reactive astrocyte signatures (IRAS1- enriched for MYC and mTORC signaling, IRAS2- enriched for IFN response genes, Leng et al. 2023).

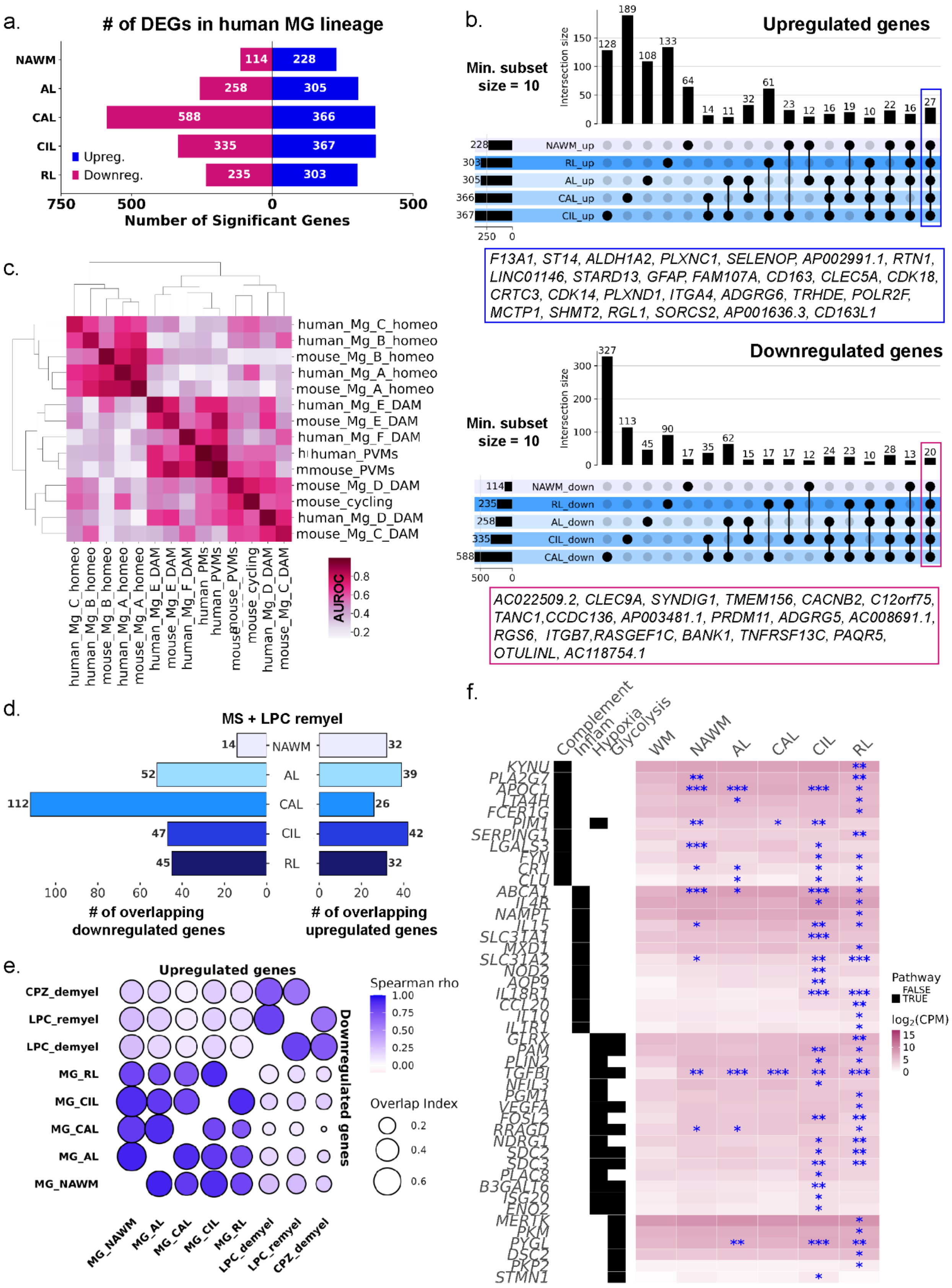

a.
### of DEGs in human MG lineage
NAWM
AL
CAL
CIL
RL
114
228
258
305
588
366
335
367
235
303
Upreg.
Downreg.
750
500
0
500
Number of Significant Genes
b.
Upregulated genes
Min. subset size = 10
Intersection size
NAWM_up
RL_up
AL_up
CAL_up
CIL_up
F13A1, ST14, ALDH1A2, PLXNC1, SELENOP, AP002991.1, RTN1, LINC01146, STARD13, GFAP, FAM107A, CD163, CLEC5A, CDK18, CRTC3, CDK14, PLXND1, ITGA4, ADGRG6, TRHDE, POLR2F, MCTP1, SHMT2, RGL1, SORCS2, AP001636.3, CD163L1
Downregulated genes
Min. subset size = 10
Intersection size
NAWM_down
RL_down
AL_down
CIL_down
CAL_down
AC022509.2, CLEC9A, SYNDIG1, TMEM156, CACNB2, C12orf75, TANC1,CCDC136, AP003481.1, PRDM11, ADGRG5, AC008691.1, RGS6, ITGB7,RASGEF1C, BANK1, TNFRSF13C, PAQR5, OTULINL, AC118754.1
c.
human_Mg_C_homeo
human_Mg_B_homeo
mouse_Mg_B_homeo
human_Mg_A_homeo
mouse_Mg_A_homeo
human_Mg_E_DAM
mouse_Mg_E_DAM
human_Mg_F_DAM
human_PVMs
mouse_PVMs
mouse_Mg_D_DAM
mouse_cycling
human_Mg_D_DAM
mouse_Mg_C_DAM
AUROC
0.8
0.6
0.4
0.2
d.
MS + LPC remyel
NAWM
AL
CAL
CIL
RL
### of overlapping downregulated genes
### of overlapping upregulated genes
e.
Upregulated genes
Downregulated genes
CPZ_demyel
LPC_remyel
LPC_demyel
MG_RL
MG_CIL
MG_CAL
MG_AL
MG_NAWM
Spearman rho
Overlap Index
f.
Complement
Inflam
Hypoxia
Glycolysis
WM
NAWM
AL
CAL
CIL
RL
KYNU
PLA2G7
APOC1
LTA4H
FCER1G
PIM1
SERPING1
LGALS3
FYN
CR1
CLU
ABCA1
IL4R
NAMPT
IL15
SLC31A1
MXD1
SLC31A2
NOD2
AQP9
IL18R1
CCL20
IL10
IL1R1
GLRX
PAM
PLIN2
TGFBI
NFIL3
PGM1
VEGFA
FOSL2
RRAGD
NDRG1
SDC2
SDC3
PLAC8
B3GALT6
ISG20
ENO2
MERTK
PKM
PYGL
DSC2
PKP2
STMN1
Pathway
FALSE
TRUE
log2(CPM)
15
10
5
0

**Supplementary Figure 12: Human microglia lineage DEG analysis.**

a. Stacked bar chart showing the total number of upregulated and downregulated differentially expressed genes in MG per lesion condition compared to WM.
b. Upset plots showing the unique (single points) and overlapping (connected points) DEGs in MG across lesion conditions that are upregulated (logFC ≥ 1.5, padj < 0.05, top) and downregulated ((logFC ≤ 1.5, padj < 0.05, bottom). Overlap size was filtered to include only points with 10 or more shared genes.
c. Heatmap of AUROC scores between mouse and human MG lineage cells. Dendrograms were generated by hierarchical clustering of Euclidean distances using average linkage.
d. Bar chart showing the overlap of upregulated and downregulated DEGs shared between MG at the LPC remyelination timepoint and the MS lesions.
e. Scatterplot matrix showing the pairwise similarity between datasets, with upper-left triangles showing concordance for upregulated DEGs and lower-right triangles for downregulated DEGs. Circle size indicates the fraction of shared DEGs (overlap coefficient). Fill color represents the Spearman correlation coefficient (ρ) of log fold-changes among shared genes and black outlines denote statistically significant correlations (p < 0.05).
f. Heatmap showing expression of genes in the indicated pathways across lesion conditions. Heatmaps are colored by the average $\log_2$-transformed CPM expression values. Gene–pathway associations were assigned to pathways using the Hallmark sets and are shown in the left annotation panel, where black tiles indicate pathway membership. Genes are hierarchically ordered first by pathway inclusion and then by average expression across all conditions. Blue asterisks indicate genes with significant upregulation (* for $p < 0.05$, ** for $p < 0.01$, *** for $p < 0.001$). (Inflam: Inflammatory Response).
